# *Helicobacter pylori* modulates heptose metabolite biosynthesis and heptose-dependent innate immune host cell activation by multiple mechanisms

**DOI:** 10.1101/2023.03.17.531716

**Authors:** Martina Hauke, Felix Metz, Johanna Rapp, Larissa Faass, Simon Bats, Sandra Radziej, Hannes Link, Wolfgang Eisenreich, Christine Josenhans

**Affiliations:** Max von Pettenkofer Institute, Ludwig Maximilians University Munich, Pettenkoferstr. 9a, 80336 Munich, Germany; Bavarian NMR Center-Structural Membrane Biochemistry, Department of Chemistry, Technical University Munich, Lichtenbergstr. 4, 85747 Garching, Germany; University Tübingen, Bacterial Metabolomics, CMFI, Auf der Morgenstelle 24, 72076 Tübingen, Germany

## Abstract

Heptose metabolites including ADP-heptose are involved in bacterial lipopolysaccharide and cell envelope biosynthesis. Recently, heptoses were also identified to have potent pro-inflammatory activity on human cells as novel microbe-associated molecular patterns. The gastric pathogenic bacterium *Helicobacter pylori* produces heptose metabolites which it transports into human cells through its Cag type 4 secretion system. Using *H. pylori* as a model, we have addressed the question, how pro-inflammatory ADP-heptose biosynthesis can be regulated by the bacteria. We have characterized the inter-strain variability and regulation of heptose biosynthesis genes and the modulation of heptose metabolite production by *H. pylori*, which impact cell-autonomous pro-inflammatory human cell activation. HldE, a central enzyme of heptose metabolite biosynthesis, showed strong sequence variability between strains, and was also strain-variably expressed. Transcript amounts of genes in the *hldE* gene cluster displayed intra-strain and inter-strain differences, were modulated by host cell contact and the presence of the *cag* pathogenicity island, and were affected by carbon starvation regulator A (CsrA). We reconstituted four steps of the *H. pylori* LPS heptose biosynthetic pathway *in vitro* using recombinant purified GmhA, HldE and GmhB proteins. On the basis of one- and two-dimensional NMR spectroscopy and mass spectrometry, the structures of major reaction products were identified as β-D-ADP-heptose and β-heptose-1-monophosphate. A pro-inflammatory heptose-monophosphate variant was also identified for the first time as a novel cell-active product in *H. pylori* bacteria. Separate purified HldE subdomains and variant HldE allowed to uncover additional strain variation in generating heptose metabolites.

## Introduction

Heptose derivatives including ADP-heptose are produced by Gram-negative and Gram-positive bacteria as building blocks for the biosynthesis of lipopolysaccharide (LPS) structures and other envelope components, such as surface layers (1). In particular, Gram-negative bacteria require heptose sugars in the inner core of their LPS. Recently, (ADP)-heptose metabolites were also defined as a class of novel microbe-associated molecular patterns (MAMPs) produced by Gram-negative bacteria. *Via* pattern recognition, these heptose metabolites, in particular ADP-heptose, can be recognized inside mammalian cells and lead to downstream NF-κB activation through the ALPK1-TIFA axis and the formation of tumor necrosis factor receptor-associated factor (TRAF)-interacting proteins with forkhead-associated domain complexes (TIFAsomes) (2–4). This innate cell activation mechanism was already described for a number of pathogenic bacteria, including *Neisseria gonorrhoeae* (5, 6), *Yersinia enterocolitica* (4), diverse *Escherichia coli* strains (4), *Shigella flexneri* (2), *Campylobacter jejuni* (7), and *Helicobacter pylori* (3, 8–12). While some bacteria release the heptose metabolites directly into the medium (7, 13), other species utilize the targeted properties of complex bacterial membrane secretion systems to inject some of these metabolites into the cytoplasm of mammalian host cells. For example, *H. pylori* appears to use mainly its Cag Type 4 Secretion System (T4SS) to also mediate the transfer of heptoses into human gastric epithelial cells (3, 10, 11) and monocyte/macrophages (12), while *Enterobacteriaceae* (*Shigella*, *Yersinia*) and *Pseudomonas* spp. may also enlist their Type 3 secretion systems (T3SS) for a similar purpose (14). Hence, similarly as for T3SS, which can transport both proteins and metabolites for host cell targeting, this also seems to be true for T4SS.

The core heptose biosynthesis pathway leading to primary production of heptoses is present in most Gram-negative bacteria and also exists in some Gram-positive bacteria (15, 16). Several Gram-negative bacteria use heptoses in their LPS inner core and in their LPS outer core or outer chains (O-antigen chains) (1, 17–19). Some bacteria incorporate heptose sugars also into their surface layers or outer polysaccharide capsules (1, 16, 20–24) and into bacterial surface-associated proteins (25). Glycosylation with glycero-manno-heptose was, for instance, detected in the AIDA-I outer membrane-associated autotransporter adhesin of intestinal pathogenic enteroaggregative *E. coli* (25), where it probably contributes to protein stabilization under harsh environmental conditions. The canonical biosynthesis pathway of LPS core heptoses leading to nucleotide (ADP)-activated heptose has first been reported in *E. coli*, *Haemophilus influenzae* and *Aneurinibacillus thermoaerophilus* (15, 16, 26–28). GDP-heptose, in addition to ADP-heptose metabolites, has also been identified, for instance in *Mycobacterium tuberculosis* and enteropathogenic *Yersinia enterocolitica* (29, 30). The canonical heptose biosynthesis pathway generally consists of five steps, which are mediated by a consecutive action of four or five different enzymes, for instance in *E. coli* and *H. pylori* GmhA, HldE (RfaE), GmhB and HldD (RfaD) (16, 31, 32). ADP-activated heptoses, delivered by the last biosynthesis steps, then can serve as substrates for the downstream glycosyltransferases involved, for example, in the assembly of the LPS inner-core oligosaccharides (33, 34).

Despite the long-standing knowledge about the canonical heptose biosynthesis pathway, very little is known about the regulation of the biosynthetic gene cluster, the diverse enzymatic activity profiles of the contributing enzymes, the role of various heptose biosynthesis genes in different bacteria, and of strain variability of those traits in different bacterial species. The gene cluster and single gene or enzyme activities have been characterized most intensively in *E. coli* (15, 28, 31, 32, 35). *H. pylori*, the model organism which is studied here, contains short heptane chains with α-configured D-glycero-D-manno-heptose units in its LPS core (19, 36). In addition, some, preferentially non-Asian, strains assemble branched heptose subunits in their outer core (19). *H. pylori* harbors one single canonical heptose biosynthesis gene cluster, consisting of *gmhA/rfaA* (HP0857), a bifunctional bi-domain-encoding *hldE/rfaE* gene (HP0858), as in *E. coli*, a *gmhB* (HP0860), and a *hldD* gene (HP0859). In *H. pylori*, the individual enzymes encoded in the core heptose biosynthesis cluster were putatively assigned and the enzymatic activities of these proteins were partially reported (37, 8). D-glycero-β-D-manno-heptose-1,7-bisphosphate (β-HBP), a major reaction product in *Neisseria* (5), and ADP-D-glycero-β-D-manno-heptose (ADP-heptose), produced by *H. pylori*, *Y. enterocolitica*, *Salmonella* and *E. coli*, were shown to exhibit an innate activation potential towards human cells via the ALKP1-TIFA axis (4, 8).

In the case of guided host cell activation by the secretion system of pathogenic bacteria, for example by transport or translocation of heptose metabolites into human cells, the important question arises, whether and how the bacteria have developed ways to cell contact-dependently and strain-dependently control the biosynthesis of phospho-heptose derivatives and their innate activity on the target cells. In order to address these open questions, we have studied *H. pylori,* which directs heptose metabolites into human cells mainly by means of its T4SS (3, 11, 12). First of all, we tested the regulation of genes in the heptose biosynthesis cluster in various strains and assessed the influence of several physiological parameters on the regulation of heptose biosynthesis in *H. pylori*. Using this approach, we have identified that strain identity, the presence of a functional CagT4SS, or the allelic diversity of the biosynthesis genes themselves contributed to strain- and condition-dependent modulation of heptose biosynthesis and metabolite activity on host cells. By reconstituting the pathway in vitro and characterizing final and intermediate products in vitro and in *H. pylori* lysates biochemically, we have also gained insight into the variation of known and novel output metabolites (products) of the heptose pathway and their activity as novel MAMPs.

## Methods

### Bacterial strains and culture conditions of H. pylori strains

*H. pylori* bacteria of various wild type strains and corresponding isogenic allelic exchange mutants in heptose biosynthesis genes, *cagY* and the *cag*PAI were used as listed with references in Table 1. The mutants were generated as reported in (11). For novel *cag*PAI deletion mutants in L7 and Su2, the insertion mutations were recreated by the same strategy as described in (11). The *cag*PAI deletion mutant is a complete deletion, while all single gene mutant were designed as insertion mutations to retain 3-prime and 5-prime segments of the genes, interrupted by a kanamycin resistance cassette. For routine culture, *H. pylori* strains were grown on blood agar plates (Oxoid blood agar plates base II) supplemented with 10% horse blood (Oxoid) and the following antibiotics (all purchased from SIGMA-Aldrich, USA): amphotericin B (4 mg/L), polymyxin B (2,500 U/L), vancomycin (10 mg/L), trimethoprim (5 mg/L), and kanamycin (optional; 10 mg/L) for selective growth of insertion mutants. For cell co-incubations, transcriptome analyses, and preparations of lysates, bacteria were freshly grown for 20 to 24 h on plates in anaerobic jars supplemented with Anaerocult C sachets (Merck, USA) at 37°C. A list of strains used in the present study is provided in Table 1.

**Table 1:** List of strains used in the present study ^a^. ^a^ all wild type strains listed are minimally passaged clinical isolates

| Strain name | Origin | Description | Reference |
| --- | --- | --- | --- |
| *H. pylori* (HP) N6 wt | hpEurope; site of isolation = France | Wild type strain, *cagPAI* positive | (55) |
| HP N6 *cagY* (HP0527) | hpEurope | Allelic exchange insertion mutant HP0527 (*cagY*) in strain N6 | (11) |
| HP N6 *hldE* (HP0858) | hpEurope | Allelic exchange-insertion mutant HP0858 (*hldE*) in strain N6 | (11) |
| HP N6 *hldE comp* | hpEurope | HP0858/*hldE* gene complementation in *rdxA* locus of HP0858 mutant in strain N6 | (11) |
| HP N6 *csrA::aphA3'-III* | hpEurope | *csrA* (HP1442) allelic exchange insertion mutant (methods) in N6 | This study, generated according to (46) |
| HP 26695a | hpEurope; site of isolation = USA | Wild type strain, *cagPAI* positive | (56) |
| HP 26695a Δ*cagPAI* | hpEurope | *cagPAI* complete deletion by allelic exchange in strain HP 26695a | (11) |
| HP 26695a *cagY* | hpEurope | Allelic exchange insertion mutant HP0527 (*cagY*) in strain HP 26695a | This study, according to (11) |
| HP L7 wt | hpAsia2, South Asia, India | Wild type strain, *cagPAI*-positive | (45) |
| HP L7 Δ*cagPAI* | hpAsia2, South Asia, India | *cagPAI* complete deletion by allelic exchange in strain L7 | This study |
| HP J99 wt | hpAfrica1, North America | Wild type strain, *cagPAI*-positive | (57, 58) |
| HP Africa2 wt | Africa2 | Wild type strain, primary *cagPAI*-negative | (57) |
| HP Su2 wt | hpNEAfrica, Northeast Africa, Sudan | Wild type strain, *cagPAI*-positive | (45) |
| HP Su2 Δ*cagPAI* | hpNEAfrica, Northeast Africa, Sudan | *cagPAI* complete deletion by allelic exchange in strain Su2 | This study |
| HP P12 wt | hpEurope | Wild type strain, *cagPAI*-positive | (59) |
| *E. coli* Rosetta pLysS | Novagen | *E. coli* protein expression strain | Novagen |

### H. pylori csrA *mutant*

The *H. pylori csrA* mutant was generated by inserting a kanamycin cassette (aphA3’-III) into the *H. pylori csrA* gene, PCR-cloned into pUC18 using *csrA-*specific primers (Table 4). The construction of the *csrA* mutant was performed with initial amplification of *csrA* (HP1442) and *csrA*-flanking regions from *H. pylori* 26695 genomic DNA and introducing PstI and KpnI restriction sites (resulting plasmid pCJ2008). Primers HP_1442del_bglII_fw and HP_1442del_bglII_rv (Table 4) were then used to reverse-amplify and religate plasmid pCJ2009 from pCJ2008. pCJ2009 contains the restriction site BglII in the center of *csrA*, with a 42 bp partial deletion of the gene. Using the BglII site, the kanamycin cassette (cut BamHI) was inserted, generating plasmid pCJ2010. The resistance cassette insertion in the chromosomal *csrA* locus of *H. pylori* strain N6 was generated after natural transformation of pCJ2010 by allelic exchange mutagenesis. *csrA* insertion mutants were selected on blood agar containing kanamycin and checked for correct chromosomal insertion-recombination using cloning primers HP_1442_PstI_fw and HP_1442_KpnI_rv as well as resistance cassette-specific primers.

### Cultivation of human cells

The human cell lines AGS (ATCC CRL-1739, human gastric adenocarcinoma cell line) and MKN28 (JCRB0253, human gastric carcinoma cell line) were routinely cultured in RPMI 1640 medium buffered with 20 mM Hepes and GlutaMAX stable amino acids (Gibco, Thermo Fisher Scientific, USA). Medium was supplemented with 10% FCS (PromoCell, Germany). AGS and MKN28 cells were cultured without antibiotics. The cell line HEK-NF-κB_luc (BPS Bioscience, USA, luciferase reporter cell line) was routinely cultured in DMEM [buffered with 20 mM Hepes] and supplemented with Glutamax (Gibco, Thermo Fisher Scientific, USA) and 10% FCS (PromoCell, Germany). HEK-NF-κB_luc were supplemented for routine growth with 50 µg/mL Hygromycin B (Invivogen, USA) in the same culture medium. For infection and coincubation experiments, all cell lines were seeded in media without antibiotics. Antibiotics were removed from all cell cultures by washing before starting co-incubation assays. All cell cultures were grown in a 5% CO_2_ atmosphere incubator and routinely passaged using 0.05% buffered Trypsin-EDTA (Gibco, Thermo Fisher Scientific, USA).

### Co-culture of cells with live bacteria or bacterial products

Human cells were co-cultured with various different *H. pylori* strains or bacterial lysates. The infection was carried out in 6-, 24- or 96-well plates (Greiner Bio-One, Austria) on sub-confluent cell layers (60%-80% confluency), seeded on the previous day. 60 min prior to infection, the medium was exchanged to fresh RPMI 1640 (supplemented with 20 mM Hepes, GlutaMAX and 10% FCS) or fresh DMEM. All bacteria-cell or compound- and lysate-cell co-incubation experiments were performed in the absence of antibiotics. For infection of cells with bacteria, *H. pylori* was harvested after 20 h of growth from fresh blood plates into cell culture medium. The OD_600_ of this bacterial suspension was measured and adjusted to the respective multiplicities of infections (MOI; MOI of 25 was used in most experiments), as indicated in the results and figures. The co-incubation was synchronized by centrifugation of the cell culture plates (300 rpm, 5 min, at room temperature). For co-incubation with enzymatically treated lysates (ETLs) or β-D-ADP-heptose (Invivogen), cell medium was also changed 60 min prior to the addition of lysate preparation or β-D-ADP-heptose to the cells to fresh medium without antibiotics, and co-incubation started with centrifugation of the plate. Co-incubation was performed in a 5% CO_2_ atmosphere cell incubator for different incubation periods (indicated in text and figures). Samples were harvested after taking off the supernatant carefully (for ELISA, IL-8 secretion), either by scraping the cells from the bottom of the plate (for RNA isolation), or by adding luminescence substrate to the cells and medium (for NF-κB luciferase quantitation). If not otherwise indicated, cells were co-incubated with bacteria or metabolite preparations for 4 h, both for measuring IL-8 secretion or performing luminescence reporter assays.

### RNA isolation of human and bacterial samples

RNA was isolated from human cells, bacteria or bacteria-cell co-incubation samples. Human cells ( co-cultured) were harvested from 6-well plates (Greiner Bio-One, Austria) or small petri dishes (diameter of 3 cm) by scraping the cells from the surface using a rubber policeman. *H. pylori* strains were harvested by resuspending bacteria from plates after approximately 20 h of growth. Human co-incubated or bacterial samples were centrifuged and pellets were snap-frozen immediately in liquid nitrogen or on dry ice and stored at −80° until RNA preparation. From pellets, total RNA was isolated using the RNeasy Mini Kit (Qiagen, Germany) following the manufacturer’s instructions after mechanical lysis of the samples in a Fastprep bead-beater (MP Biomedicals Inc., USA), at 5 MHz for 45 s using lysing matrix B (for bacterial samples; MP Biomedicals). Isolated RNA was DNase-treated using TURBO DNAse clean-up kit (Ambion-Invitrogen, USA). Sufficient RNA quality and purity were ensured by photometric measurement, gel electrophoresis and RNA ScreenTape analysis in a Tape Station (Agilent, USA), using high sensitivity RNA tapes, and by the amplification of control genes (*H. pylori* 16S rDNA for control of RNA purity). Total RNA was then used for genome-wide RNA-sequencing (RNA-Seq), or further processed for cDNA generation and performance of RT-qPCR.

### cDNA synthesis and RT-qPCR

cDNA was synthesized from 1 µg of total bacterial RNA using Superscript III reverse transcriptase (Invitrogen, USA) RNase-Out (Invitrogen, USA) and random nonamer primers. Reverse transcription was performed from 1 µg of total RNA. All reagents were purchased from Invitrogen, USA. Sufficient quality of cDNA was ensured by control PCRs (amplification of 16S for bacterial cDNA).

Quantitative RT-qPCR was routinely performed on 0.5 µl cDNA in a CFX96 real-time PCR cycling machine (BioRad, USA) using *H. pylori* gene-specific primers (synthesized at Metabion, Table 2), and 2x SYBR Green Master Mix (QIAGEN, Hilden, Germany). Transcript quantification was always performed in triplicates, with gene-specific standards for absolute amount quantification, performed in parallel for each transcript, using the following protocol: 95° 10:00, (95° 0:30, 55° 0:30, 72° 0:30, 40 cycles), melt curve 60° to 95°, increment 0.5° for 0:05. Results were equalized to 1 µL of cDNA and normalized to transcript amounts of the *H. pylori* 16S gene, using a correction factor respective to the mock reference, but maintaining the absolute values (pg/µl) for all final transcript amounts. MiQE standards for the qPCR method were applied as referenced in (37). Each figure shows different biological experiments, which explains the slight variation of absolute transcript quantities. Different experiments can have slightly different starting quantities/transcript amounts of certain transcripts, due to biological variation.

**Table 2:**
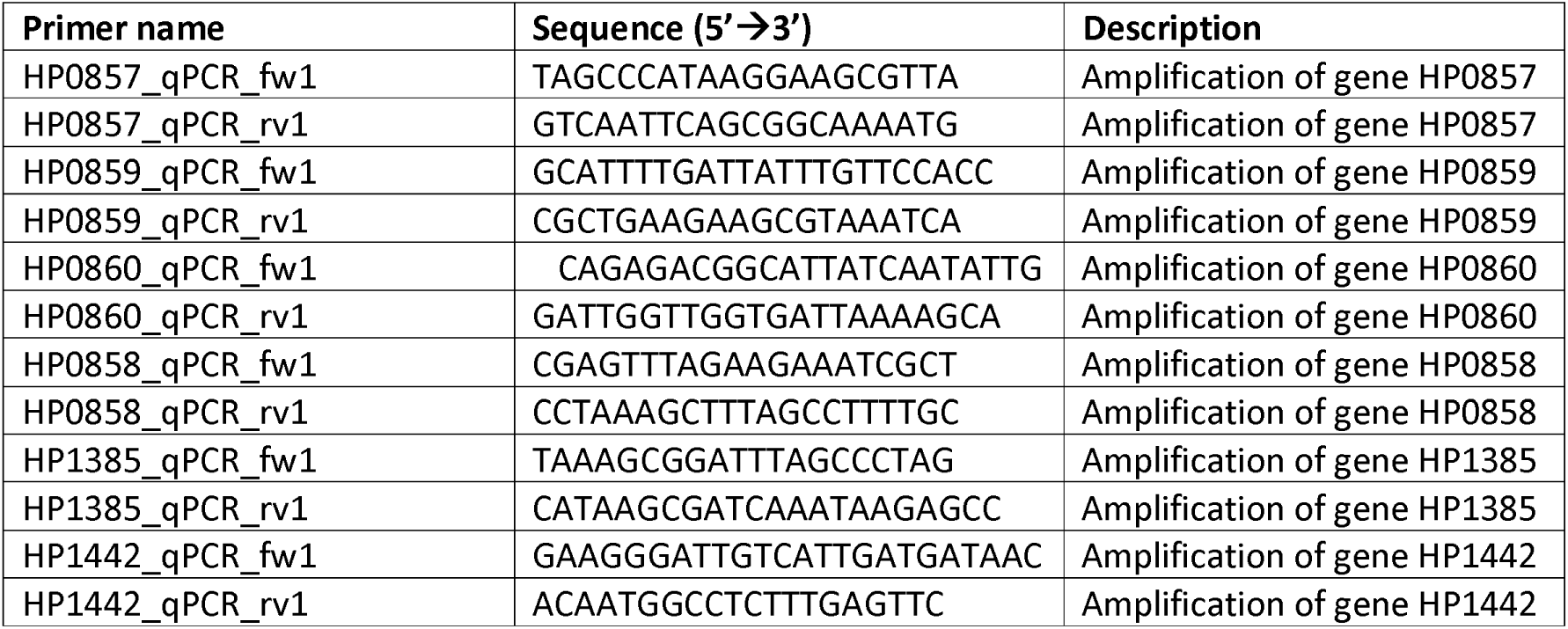
List of gene-specific primers used for qPCR.

### RNA-sequencing and data analysis

RNA-Seq and transcriptome analysis were performed as previously described (12). Briefly, paired, trimmed, quality-filtered, fastq files with an average read length of 150 bp, obtained from the Illumina NextSeq2500 platform, were processed using the CLC Genomics Workbench Version 20.0 (QIAGEN, Germany). Paired reads were down-sampled for each sample to an equal of 2,000,000 and mapped against the *H. pylori* reference genome 26695 (accession AE000511, NCBI database; (38)). Reads were aligned as previously described (12) and gene expression levels were quantitated as RPKM values, normalized to the gene length and read number. RPKM values were then used for calculating differential expressions between samples using the CLC Genomic Workbench RNA-Seq analysis workflow. See Table S2 for full transcriptome results and differentially expressed genes. Full results are accessible on NCBI GEO (gene expression omnibus) under accession number GSE227450.

### ETLs (enzymatically treated lysates) and water lysates

Bacterial lysates were prepared from *H. pylori* harvested from blood agar plates after 20 h of growth into sterile cell culture PBS (1x concentration; Gibco, Thermo Fisher Scientific, USA). The bacterial suspension quantified by OD_600_ measurement was centrifuged and the pellets were stored at −20°C. Briefly, for lysate preparation, the frozen pellet was resuspended in an appropriate volume (1 ml) of 1 x PBS and adjusted to an OD_600_ of 2 (per mL). The suspension was boiled for 10 min in a water bath (99°C) and centrifuged. The supernatant was sterile-filtered (0.22 µm pore filter, Merck Millipore, Germany) and stored at −20°C if not further processed immediately. Further processing of the samples was performed either enzymatically (generation of ETLs, as in: (11) or by methanol precipitation (one part bacterial preparation and three parts methanol). Highly purified ETLs were prepared, treating the bacterial lysates as previously described (11). For NMR measurements (less-pure lysate preparations, which still contain, for instance, residual DNA), lysates of an OD_600_ of 20, 10 or 5 in one ml were prepared as described and boiled for 20 min. These lysates were then additionally treated for protein precipitation with 3 volumes of ultra-pure methanol (Sigma-Aldrich, USA). Ultimately, samples were vortexed and centrifuged to precipitate proteins and to recover protein-free supernatants, which were then sterile-filtered as described above. ETL samples used for mass spectrometry were additionally mixed with methanol and acetonitrile (1:2:2 (v/v/v) final mixture).

### Cytokine measurement from cell supernatants

IL-8 secretion of human cells into supernatant was quantified performing human IL-8 ELISA, according to the manufacturer’s protocol (BD OptEIA #555244) on pre-tested sample dilutions. Colorimetric signal detection was performed using a Clariostar multiwell plate-reading machine (BMG labtech).

### Luciferase quantitation in human reporter cells

Briefly, firefly luciferase signal of HEK-NF-κB_luc cells was determined using Steady Glo/Bright Glo Luciferase Assay (Promega, USA), as previously described (12). Luciferase quantification was regularly performed in 96-well-F-bottom plates (Greiner BioOne, Austria, # 655180) and using 50 µL total sample volume. After 4 h of co-incubation with live bacteria, bacterial lysates, β-D-ADP-heptose, or other metabolites as indicated in results and figures, an equal volume of the luciferase lysing and detection buffer (Promega) was added to each well. The reaction was allowed to incubate for 10 min for cell lysis with shaking, followed by luminescence measurement in a Victor Nivo Multimode Microplate Reader (PerkinElmer, USA) at the following settings: shaking for 3 s, no filter, 1 s photon counting. All conditions were analyzed in duplicates or triplicates.

### SDS-PAGE and Western Blot for protein detection and quantification

Bacterial samples were prepared by harvesting bacteria directly from the plate, after centrifugation, into 1 x PBS, followed by ultrasonication (Branson sonifier, 2 x 1 min at power setting 5), and subsequent separation of soluble and insoluble fractions by centrifugation (10,000 x g, 20 min, 4°C). Protein concentration of samples was determined by BCA assay (Pierce, Thermo Fisher Scientific, USA) using a Clariostar multiwell reader for final colorimetric reading. Regularly, 10 µg of protein, equalized amounts for all samples, was loaded onto 11.8-14% SDS gels, and run at 100 V constant voltage in Laemmli buffer supplemented with 0.1% SDS. Blotting was performed onto BA85 nitrocellulose membranes (Schleicher & Schuell) in Towbin buffer for 2 h at 300 mA. Blotted membranes were blocked using 5% skim milk (Sigma-Aldrich, USA or BioRad, USA), or 1 to 5% BSA in TBS buffer containing 0.1% Tween (TBS-T), according to antibody specifications. Specific antibody (Table 3) incubations were performed for 1 h at ambient temperature or incubated overnight at 4°C. As secondary antibody, goat-anti-rabbit or goat-anti-mouse antibody, coupled to horseradish peroxidase (Jackson Immuno Laboratories) was used at a dilution of 1:10,000. Signal was detected using Immobilon HRP chemiluminescence substrate detection reagent (Merck Millipore, Germany) and imaged in a chemiluminescent imager (BioRad, USA).

**Table 3:** Specific antisera and antibodies used in this study.

| Antigen | Source species | Reference |
| --- | --- | --- |
| HIdE | Rabbit | This study |
| FlhA | Rabbit | (60) |

### DNA cloning methods for protein expression

For expression of heptose biosynthesis enzymes, genes HP0857 (*gmhA*), HP0858 (*hldE*) and HP0860 (*gmhB*) were cloned from PCR products generated from genomic DNAs of the respective *H. pylori* wt strains, 26695a and N6, using BamHI and NotI (NEB, USA) as full-length constructs in pET28a(+) (EMD Biosciences, Novagen, Germany). The two strains were selected for their diversity in *hldE* sequence. To express the two domains of the bifunctional enzyme HldE (HP0858), the first (amino acids 1-325) and last segments of the gene (amino acids 328-461) were cloned separately into pET28a(+) using the same restriction enzymes. Gene constructs were located behind a 6x N-terminal His-tag, separable from the enzyme using the TEV protease cleavage site between construct and His-tag. The plasmids confer kanamycin resistance allowing selection of clones. Clones were checked by restriction analysis and Sanger sequencing of the complete inserts. Resulting plasmids: HP0858/*hldE* [strain 26695] is pCJ1628; HP0858/*hldE* [strain N6] is pCJ2004; HP0857/*gmhA* [26695] is pCJ1627; HP0860/*gmhB* [26695] is pCJ1630; HP0858/*hldE* [26695] d1 = pCJ2002; HP0858/*hldE* d2 = pCJ2003. The primers used for cloning are listed in Table 4.

**Table 4:**
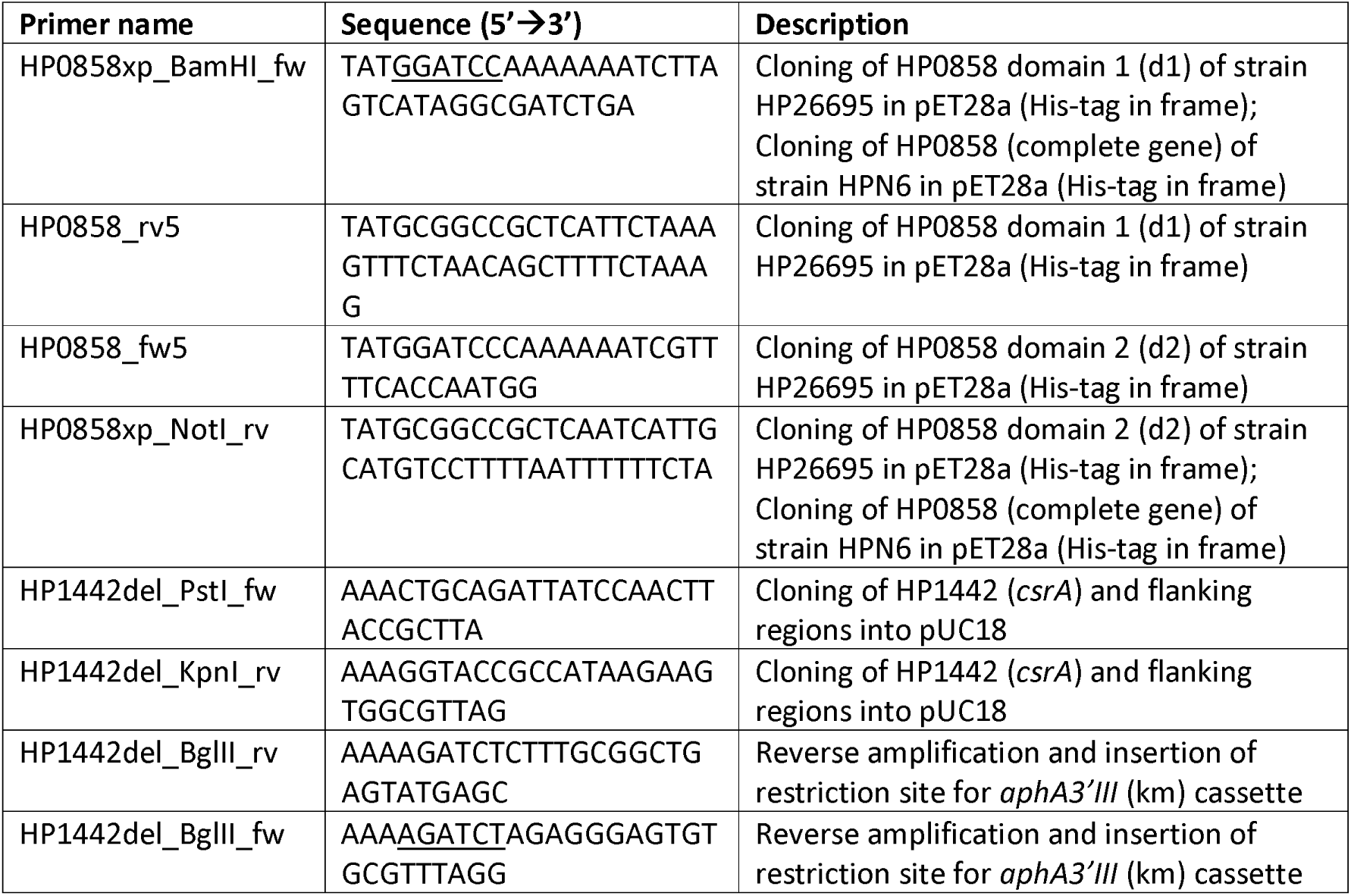
Primers for cloning.

### HEK cell transfection for metabolite activity testing (ETLs)

HEK-NF-κB_luc reporter cells (BPS Bioscience, USA) were transfected with bacterial ETLs or *in vitro*-synthesized heptose metabolites using Lipofectamine 2000 transfection agent (Invitrogen). Transfection was performed in 96-well plates, and cells were seeded to 3×10^4^ cells in 50 µL DMEM (containing 10% FCS) per well, approximately 20 h prior to transfection. Medium was exchanged to 25 µL OptiMEM (containing 5% FCS), 1 h before addition of 25 µL transfection agent containing 4 µL ETL or heptose metabolites, 0.5 µL Lipofectamine 2000, and 20.5 µL OptiMEM per well. Transfected cells were subsequently incubated at standard culture conditions; after 4 h incubation, luciferase substrate buffer (SteadyGlo, Promega) was added for cell lysis and luminescence was detected as described above.

### Protein expression and purification from E. coli

Expression of heptose biosynthesis enzymes from inducible expression plasmids was carried out in Luria-Bertani broth (LB, Lennox [Oxoid]) medium supplemented with 50 µg/µL kanamycin (Sigma, USA), from expression cultures set up in *E. coli* Rosetta pLysS (Novagen/Merck, Germany), inoculated from overnight culture to a starting OD_600_ of 0.1. At an OD_600_ of 0.5 to 0.8, expression cultures were induced with 0.1 to 0.5 mM IPTG and grown to express protein for 4.5 h at 30° (with 175 rpm shaking). Bacterial pellets were harvested by centrifugation (10,000 x g, 10 min, 4°C) and stored at −20°C prior to further purification. Proteins were purified from the soluble (native) fraction, after the lysis of bacterial pellets by sonication, using a 1 ml Ni-NTA FPLC affinity chromatography column (Macherey & Nagel, Germany) coupled to an Äkta Purifier or Prime system (Cytiva/GE Healthcare, U.S.A) at 20°C (room temperature) or 4°C (the latter for N6 HldE and 26695 GmhB). After loading the column, non-specific impurities were removed by extensively washing the column with purification buffer (50 mM Tris/HCl pH = 7.5, 300 mM NaCl, 1mM DTT) and high-salt purification buffer (1 M NaCl added to the previous), followed by equilibration in purification buffer. Gradient elution was performed by increasing the imidazole concentration of the purification buffer/lysis buffer gradually from 0 to 500 mM. High protein content elution fractions were pooled, dialyzed into buffer without imidazole and further characterized and protein purity and quantity analyzed in SDS gels using appropriate reference protein loadings.

### Generation of an anti HldE antiserum

The antiserum was raised in rabbit against overexpressed, column-purified full-length *H. pylori* HldE protein (cloned from strain 26695).

### In-vitro reconstitution of H. pylori heptose biosynthesis pathway

All three central proteins of the pathway, GmhA, HldE, GmhB, cloned from *H. pylori* into pET28a vector, were overexpressed and purified from *E. coli* by Ni^2+^ affinity chromatography (see above). All proteins were purified from the soluble fraction in native form. *In vitro* reconstitution of the biosynthetic pathway was performed as follows: about 5 to 10 nmole (1-2 µg) of each protein were combined to a 50 µl reaction in an inert reaction buffer without any phosphate and with or without the substrate seduheptulose-7-phosphate (S7-P) (buffer containing: 20 mM Hepes pH = 7.5, 5.8 mM ATP, 20 mM KCl, 10 mM MgCl_2_); reactions were incubated overnight (approximately 18 h) at 37°C. After heat inactivation (HI), this material was directly used for all cell co-incubations. For NMR analysis, the samples were boiled to inactivate any proteins or enzymes and precipitated with methanol (1:3 (vol/vol); 1 volume reaction mixture, 3 volumes methanol) to remove denatured protein. The supernatant after protein removal by centrifugation, containing the metabolite reaction products, was then concentrated to dryness by a flow of gaseous nitrogen. For mass spectrometry, the reaction mixes were scaled up to 300 µl volumes.

### NMR and mass spectrometry analysis of bacterial heptose metabolites and reaction products

#### Reference ADP-heptose for NMR analysis

50 μl of each commercial reagent, β-D-ADP-heptose (2 mM, Invivogen) and β-L-ADP-heptose (J&K, China), was dried under a stream of nitrogen and each dissolved in 120 μl D_2_O. Those were used as references for the chemical identity of *H. pylori*-produced ADP-heptose isomers.

#### Sample preparation

The following samples were prepared for NMR analysis of heptose metabolites (compare also Table 6): #2: control sample containing 20 mM HEPES buffer at pH 7.5 (20 mM KCl, 10 mM MgCl2 and 6 mM NaATP) and *H. pylori* purified pathway enzymes GmhA, HldE and GmhB, but no pathway substrate; #6: HldE and substrate seduheptulose-7-phosphate (S7-P, 2 mM, Sigma-Aldrich) putative product = seduheptulose-7-phosphate (S-7-P); #11: enzymes GmhA and HldE and substrate seduheptulose-7-phosphate (2 mM, Sigma-Aldrich) - putative product = β-D-heptose-1,7-bisphosphate (d-glycero-β-d-manno-heptose 1,7-biphosphate, β-D-HBP); #9: enzymes GmhA, HldE and GmhB, and substrate seduheptulose-7-phosphate (2 mM) → putative product = ADP-heptose; All samples were incubated for metabolite biosynthesis at 37°C overnight and subsequently heat-inactivated at 95°C for 20 min. Afterwards, all samples were treated with MeOH (3:1 MeOH: samples) for protein precipitation and pelleted by centrifugation. Each of the supernatants was dried under a stream of N_2_ at about 40 °C for 30 min, and the remaining residue was dissolved in 120 μl D2O. Each of the solutions was carefully filled into a Bruker NMR Match micro tube.

#### NMR spectroscopy for metabolite identification and characterization

All NMR spectra were recorded with a Bruker AVANCE 500 MHz spectrometer equipped with a SEI probe using TopSpin Version 3.5 (Bruker Biospin GmbH, Rheinstetten, Germany). ^1^H NMR spectra were recorded with the Bruker pulse program “noesygppr1d” for suppression of the water signal during the relaxation period, applying a narrow saturation pulse with a line width of about 25 Hz. The parameters were ns = 64 or 256, ds = 8 s, TE = 25°C, aq = 2.73 s, td = 32768, sw = 12.0 ppm, and p1 = 8.20 μs. ^1^H,^1^H-COSY spectra with water suppression were recorded with the Bruker pulse program “cosygpprqf” with ns = 4 or 64, ds = 8 s, TE = 25°C, aq = 0.10 s, td = 1024 (f2), 256 (f1), sw = 10.00 ppm, and p1 = 8.20 μs. Data were processed with MestreNova Version 14.2.0 (Mestrelab Research, Santiago de Compostela, Spain). The FIDs were zero-filled and multiplied by a mild Gaussian function prior to Fourier transformation. Under these conditions, the detection limit for ADP-heptose was approximately 27 µg of dissolved ADP-heptose, i.e. 361.8 µM (applying 128 scans in the one-dimensional experiment). The ^1^H-NMR spectrum of 830 µM ADP-heptose is shown in Figure 1. With the help of ^1^H,^1^H-COSY and by comparing to previously published data (40, 41), the detected signals could be clearly assigned to protons of the adenine unit, the ribose unit and the heptose unit. Further on, characteristic CH_3_-signals at 1.21 ppm and CH_2_-signals at 3.14 ppm belonging to trimethylamine were detected for the reference compound. Notably, triethylamine is often used during the chemical synthesis of ADP-heptose as a protection reagent and is present as the counter ion in the reference sample (8). Chemical shifts, multiplicity, coupling constants as well as correlations observed in the ^1^H,^1^H-COSY spectrum for substrates, reference compounds (S7-P, β-D-ADP-heptose, β-L-ADP-heptose) and products are available upon request.

**Fig 1.**
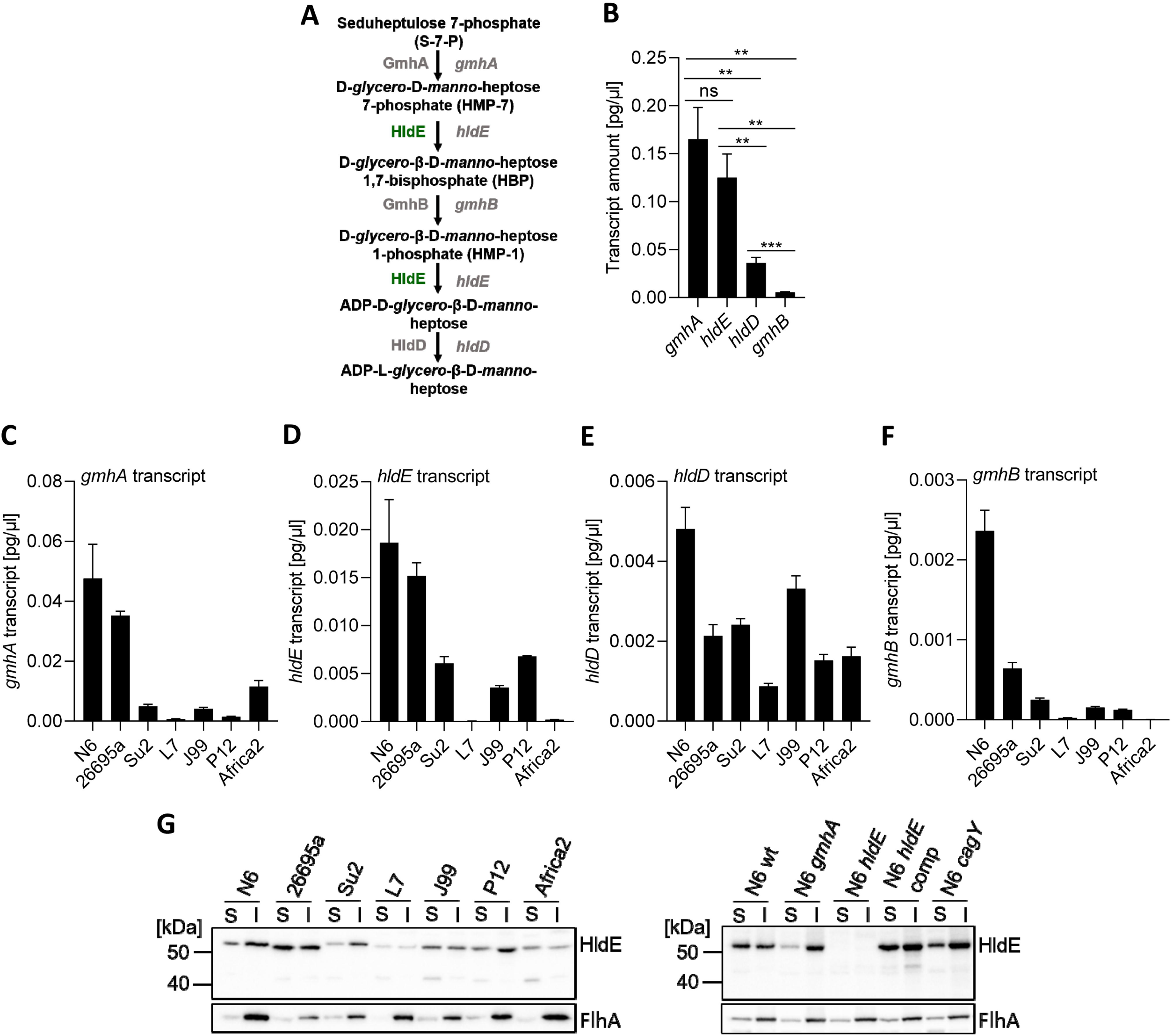
Expression of heptose biosynthesis cluster genes and gene product HldE in *H. pylori* is gene-specific and strain-specific. **A)** Schematic of the *H. pylori* 5-step ADP-L-heptose biosynthesis pathway with enzymes and proposed intermediate reaction products. The central enzyme HldE/RfaE (green) is encoded by the gene HP0858 (*hldE*) and is a bifunctional enzyme performing two steps of the synthesis cascade. **B)** Quantification of absolute transcript amounts (pg/µl) of each transcript of heptose biosynthesis cluster genes *gmhA*, *hldE* (HP0858), *hldD,* and *gmhB*, on *H. pylori* strain N6 cDNA, determined by RT-qPCR. **C) to F)** show a selection of diverse *H. pylori* wild type strains of different geographical origins was used to compare the transcript amounts of genes *gmhA* (C), *hldE* (D), (*hldD*/*rfaD*) (E), (*gmhB*) (F) between different strains. All qPCR reactions were performed in technical triplicates. The results are given as absolute values for transcript amounts (pg/µl), normalized with a correction factor to 16S rRNA transcript amounts for each strain. Strain N6 was used as a reference for normalization. **G)** Western immunoblot (anti-HldE antiserum, 1:20,000) detecting the bifunctional protein HldE and the loading and fractionation control protein FlhA (flagellar membrane protein) in soluble (S) and insoluble (I) fractions of different *H. pylori* wild type strains (left panel), and of selected isogenic mutants (right panel) of the *H. pylori* strain N6. 10 µg of total protein was loaded in each lane to provide equalized amounts. Comparisons for statistically significant differences in panel B) were performed between all conditions using unpaired student’s *t*-test; significant p values are marked with asterisks: ** p < 0.01; *** p < 0.001. Statistics for all pairwise comparisons in panels C, D), E), F) were performed by two-way ANOVA, followed by Tukey’s test, and are summarized in separate Table S1.

#### Mass spectrometry (LC-ESI MS/MS) of heptose metabolites

Pure reagents, in vitro reaction mixes or cell extracts were analyzed on an Agilent 6495 triple quadrupole mass quadrupole mass spectrometer equipped with an ESI ion source and coupled to an Agilent 1290 Infinity II UHPLC (both Agilent Technologies). Chromatographic separation of ADP-heptose was achieved with an Acquity UPLC BEH Amide (1.7 µm, 2.1×100 mm) column (Waters). Solvent A: water with ammonium formate (10 mM) and formic acid (0.1 % v/v). Solvent B: acetonitrile with formic acid (0.1% v/v). The LC-Gradient was: 0 min 90 % B, 7 min 40 % B, 8 min 40 % B, 8.5 min 90 % B, 12 min 90 % B (flow rate 0.1 mL*min-1). The heptose-phosphates were separated by a SeQuant ZIC-pHILIC (5 µm, 150×2.1 mm) column (Merck). Solvent A: water with ammonium carbonate (10 mM) and ammonium hydroxide (0.2 %). Solvent B: acetonitrile. The LC-Gradient was: 0 min 90 % B, 12 min 40 % B, 14 min 40 % B, 15 min 90 % B, 18 min 90 % B (flow rate 0.2 mL*min-1). 3 µL was injected per sample. The settings of the ESI source were: 200 °C source gas temperature, 14 L*min-1 drying gas and 24 psi nebulizer pressure. The sheath gas temperature was at 250 °C and flow at 11 L*min-1. The electrospray nozzle was set to 500 V and capillary voltage to 2500 V. ADP-heptose was analyzed in positive ion mode with a transition from 620 m/z to 428 m/z (collision energy: 10 keV, dwell time 100 ms). Heptose-bisphosphate was analyzed in positive ion mode with a transition from 371 m/z to 273 m/z (collision energy: 10 keV, dwell time 100 ms). Heptose-monophosphates were analyzed in negative ion mode with a transition from 289 m/z to 79 m/z (collision energy: 40 keV, dwell time 100 ms). Raw data were converted into text-files using MSConvert (42). Data analysis was performed with a customized Matlab script. Bioblanks were also measured by spiking the reference compounds into bacterial lysates, in order to determine possible shifts in retention times.

## Results

### Strain-specific sequence diversity, differential regulation of genes, and protein expression in the *H. pylori* LPS heptose biosynthesis gene cluster

To form a basis for assessing the heptose biosynthesis capacities in *H. pylori* and their strain-dependent genetic diversity, we initially analyzed the genomic organization of the heptose biosynthesis gene cluster and sequence polymorphisms of its cluster genes in different *H. pylori* strains. The *H. pylori* heptose biosynthesis gene cluster, similar to the *C. jejuni* cluster (7), has a counterintuitive organization, which is conserved in all *H. pylori* strains: the genes which are relevant in the later steps of the biosynthesis pathway are transcribed first and located directly downstream of the non-translated 3’ region. The *gmhA* gene (HP0857), which codes for the first enzyme in the pathway, is the last gene of the cluster (Fig. S1A). *hldE* (HP0858), the second last gene in the operon, encodes a bi-functional enzyme with two domains (d1 and d2) which are required for two separate steps in the biosynthesis pathway (Fig. 1A, Fig. S1A). Interestingly, the *H. pylori hldE* gene displays a relatively high sequence diversity between strains, while the other genes of the cluster and their predicted protein products are highly conserved (Supplemental Fig. S1B), as expected for a housekeeping function. As a basis for further functional investigations, we sought to clarify fundamental questions regarding the transcript quantities of the *H. pylori* heptose biosynthesis cluster genes and how those genes might be regulated. First, we determined the transcript amounts within the gene cluster for strain N6. This is the only strain where we could obtain insertion mutants in all heptose cluster genes (11). While the first genes in the cluster, presumably expressed from a housekeeping sigma^80^ promoter (43), had comparably low transcript amounts, the downstream genes in the cluster, *hldE* (HP0858) and *gmhA* (HP0857), displayed much higher transcript levels (Fig. 1B). This cannot be explained by the known transcript start sites identified in reference strain 26695 (44), where no additional start sites (primary or processed) are located directly upstream of HP0858 in the heptose gene cluster. Genome-wide transcriptome analyses revealed that the respective transcripts are strongly regulated by growth phase and by m^5^C DNA methylation ((38); and own unpublished data). We next addressed further internal and external influences on their gene regulation and possible regulation mechanisms.

### Expression of LPS heptose biosynthesis genes and pro-inflammatory host cell activation are *H. pylori* strain-specific

Earlier work (45) established that *cag*-positive, live *H. pylori* strains can induce very different activation levels in gastric epithelial cells at early time points, which can now be attributed almost exclusively to heptose metabolite signaling *via* the ALPK1-TIFA pathway (2, 11). Hence, we asked the question which differences in heptose biosynthesis (genetic or metabolite output) between strains might exist that can be responsible for the strain-variable activation potential.

We first tested seven different *H. pylori* strains from different geographical locations and bacterial populations (45, Table 1) for their heptose gene cluster transcripts (Fig. 1C to 1F). Strong strain-specific differences in transcript amounts for each single gene were detected, with up to two logs of differences in absolute specific transcript quantities between strains (e.g. between N6 and Su2). The highest absolute transcript quantities in most strains, but also strongest quantitative transcript differences between strains, were obtained for transcripts of *gmhA* and *hldE* (Fig. 1C, Fig 1D). In different strains, divergent patterns of relative transcript amounts of heptose gene cluster genes were determined (Supplemental Fig. S1C to S1G). According to relative transcript amount differences of *gmhA*, *hldE*, *gmhB*, and *hldE*, strains can be grouped into two major clusters (A and B), of which cluster A strains (such as: 26695a, N6, Su2) had the highest expression of both *gmhA* and *hldE* transcripts. In contrast, strains in cluster B (examples: L7 and Africa2, a primary *cag*PAI-negative isolate) exhibited the highest expression of *gmhA* and *gmhB* (HP0860), whereas the second functional gene of the pathway, *hldE*, was much less expressed (Fig. 1A, Suppl. Fig. S1A, Fig. S1C to S1G). This seems to suggest that the strains have evolved distinct patterns concerning enzymatic activities and potential metabolite output of the heptose pathway.

To back up the transcript analysis by other methods focusing on pathway output, we assessed pro-inflammatory activity of pathway products and compared them with transcript levels of the pathway genes for various strains. Enzymatically treated lysates of the bacteria (ETLs), as demonstrated previously (11), are a good proxy measure for overall content in pro-inflammatory cell-active heptose metabolites in the bacteria, when grown independently of cells. In previous work, we had only tested two different wild type strains’ ETLs (11), which appeared to have comparable pro-inflammatory MAMP activities, dependent on active heptose biosynthesis, but appeared to be significantly lower than for live bacteria. In order to clarify this preliminary result, we tested live bacteria and ETL (treated lysates) generated from seven *H. pylori* wild type strains (six *cag*PAI-positive, one primary *cag*PAI-negative) for their activity on AGS, MKN28 stomach epithelial cell lines and on HEK_luc NF-κB-luciferase reporter cells (Fig. 2, Suppl. Fig. S2). Most strains’ ETLs activated pro-inflammatory cell responses at a comparable, low level (Fig. 2A), and similar to the response to a reference amount of β-D-ADP-heptose (read-out was luciferase reporter quantification or IL-8 secretion, and the results of an ADP-heptose titration are shown in Fig. S4B). The only exception for ETL activity was strain N6, which reproducibly activated cells about two-fold more by its cleared lysate than the other strains (Fig. 2A), suggesting a higher content in pro-inflammatory metabolites. Those results indicated an overall low innate activity for preformed heptose pathway products contained in bacteria grown in the absence of cells, in particular suggested for those metabolites that can be taken up actively by the target cells (41). The live bacteria, except for the *cag*PAI-deficient strains, all activated NF-κB and elicited IL-8 secretion, with significant inter-strain differences in heptose-dependent cell activation (Fig. 2B, 2D), as already demonstrated before (44). Pro-inflammatory activation by live bacteria was about ten-fold higher than ETL-mediated activation (IL-8 or NF-κB reporter quantitation; Fig. 2). The relative pro-inflammatory activation patterns between the tested ETLs from different strains were comparable for the different epithelial cell lines used (Fig. 2B, Fig. S2). The results likewise underlined that live bacteria and the active CagT4SS provide the most important means of targeted cell transport for heptose metabolites in wild type isolates.

**Fig 2.**
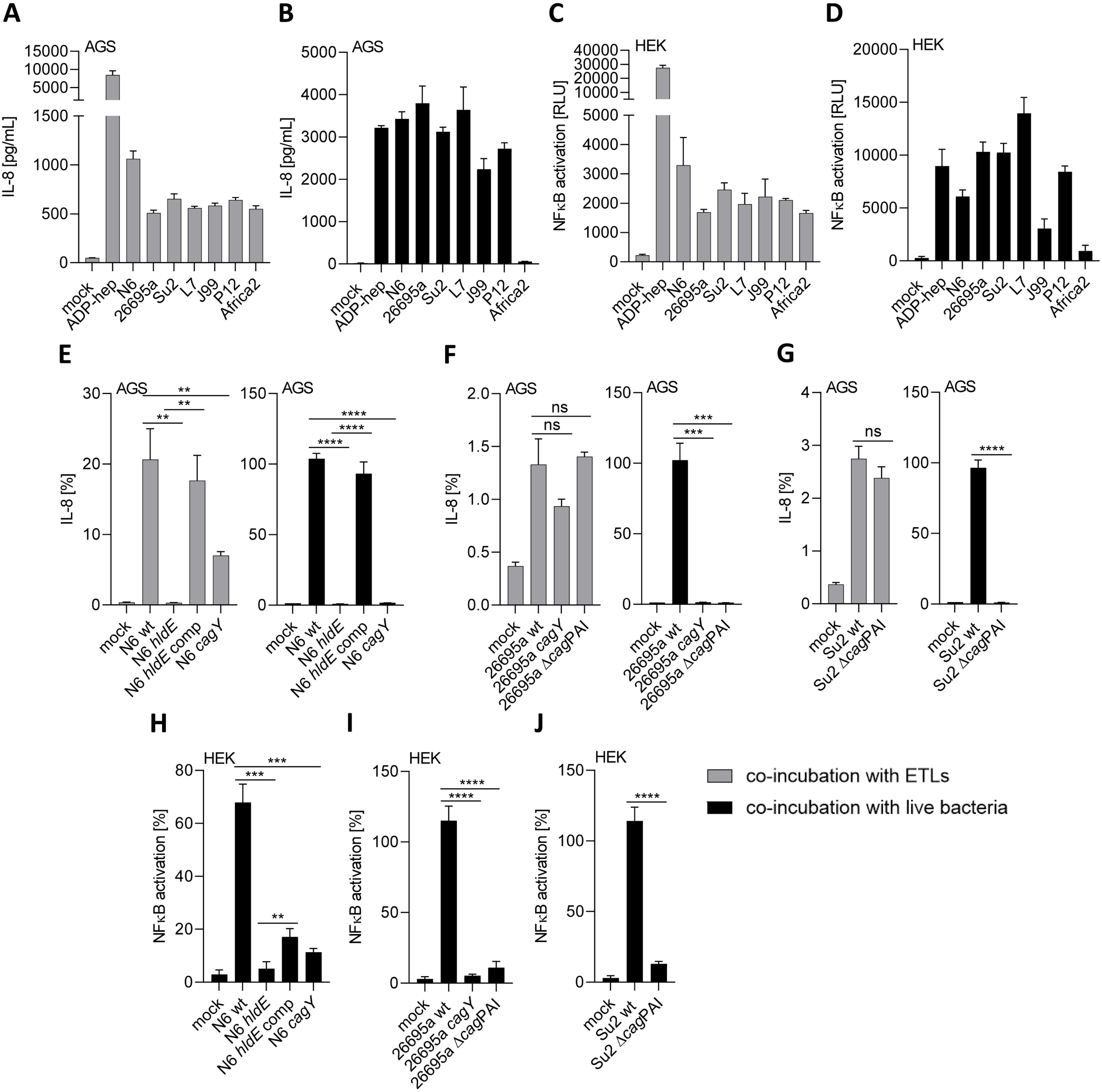
Co-incubation of epithelial cells with ETLs from various *H. pylori* strains and corresponding live bacteria demonstrates distinct potential in pro-inflammatory cell activation. **A)** through **D)** Pro-inflammatory activation of gastric epithelial AGS cells (A, B) or HEK cells (C, D) by ETLs prepared from wild type strains (A, C) or corresponding live bacteria (B, D) (MOI=25). Cells were co-incubated with ETLs or live bacteria for 4 h; activation was measured by IL-8 ELISA on cell culture supernatants (AGS) or luminescence measurement of NF-kB-luc reporter cell line (HEK), respectively. Absolute values of the response are depicted for IL-8 (in pg/ml) and for luciferase measurements in relative luminescence units (RLU) as shown on the y-axes; for comparative purposes, we applied ADP-heptose (2.5 µM) as a control stimulant for each experiment. **E)** to **G)**: activation of AGS cells after co-incubation with ETLs from different *H. pylori* wild type and mutant strains or the corresponding live *H. pylori* bacteria (MOI=25) for 4 h. In grey: Cell activation by ETLs produced from *H. pylori* wild type strains and mutants; in black: AGS cell activation by live bacteria. Cell responses were quantitated in each experiment by IL-8 ELISA from cell culture supernatants. Triplicate measurements of biological duplicate experiments are shown. Experiments were repeated at least twice with comparable results. **H)** to **J)** Pro-inflammatory cell activation of HEK-NF-κB_luc reporter cells after co-incubation with live *H. pylori* bacteria for 4 h. Cell responses were quantitated by measuring luminescence. All conditions were performed in biological triplicates and experiments were repeated at least once on different days. The results in E), F), G), H), I), and J) are depicted in percent values and were normalized to a ADP-heptose-exposed (at 2.5 µM, not shown) control condition performed in each experiment. Comparison for statistically significant differences in panels E), F), G), H), I), J) was performed between wild type-exposed sample and each mutant-exposed sample using unpaired student’s *t*-test; Significant p values are marked with asterisks: ** p < 0.01; *** p < 0.001; **** p < 0.0001; ns = non-significant.

In order to assess the role of additional bacterial functions for heptose metabolite activity, we generated another set of mutants and compared them with their isogenic wild type strains. Those included complete *cag*PAI deletion mutants (strains 26695a, Su2, and L7) (Table 1 for strain list), and in strain N6, where we could not obtain a *cag*PAI deletion, a *cagY* insertion mutant (defective T4SS). For N6, we also used the previously characterized isogenic *hldE* mutant and an *hldE*-complemented strain as heptose-negative and -positive references, respectively ((11); and Table 1). When we generated ETLs from the *H. pylori* isogenic mutants in the heptose pathway and of mutants deficient in T4SS assembly and tested them in comparison to the respective wild type strain ETLs on various cell types and reporter cell lines, we found a strong difference in epithelial cell activation (Fig. 2A–2G), which was entirely dependent upon HldE activity and influenced by an active CagT4SS. ETL from *cagY* mutants had an intermediate phenotype of pro-inflammatory cell activation on AGS cells, lower than the wild type, while ETL from Δ*cag*PAI mutants showed no change in pro-inflammatory activation compared with the wild type. Live bacteria of the same strains and mutants (Fig. 2E-2J) exhibited a different activation pattern in comparison to the ETLs. Live *hldE* mutant bacteria showed almost no cell activation (as previously published; (11)), while live bacteria from *cagY* and Δ*cag*PAI mutants (both T4SS-deficient), strain-independently, activated cells significantly less than their corresponding parental strains. This outcome confirmed that live bacteria generally require an active T4SS for strong innate immune activation of epithelial cells. We next analyzed bacterial lysates of our set of seven diverse *H. pylori* strains by Western immunoblot (Fig. 1G), using a custom-produced polyclonal antiserum against HldE, in order to detect strain-specific differences in protein expression. The serum was highly reactive against a protein of the predicted mass (ca. 65 kDa) in all tested *H. pylori* strains, but not in an *hldE* mutant (Fig 1G). The serum identified a very variable expression of HldE protein in the various assayed *H. pylori* wild type strains (Fig. 1G). We detected rather uniform HldE expression in a range of isogenic heptose (11) and *cag* mutants in *H. pylori* N6 (Fig. 1G).

### Regulation of heptose biosynthesis pathway transcripts in *H. pylori* is influenced by the absence or presence of the *cag*PAI or contact to human cells

We hypothesized that altered environmental conditions and the presence or functionality of the CagT4SS in *H. pylori* can influence the activity of the heptose biosynthesis pathway, since heptoses will be transported by the secretion system, and, possibly, the T4SS can influence production and identity of cell-active pro-inflammatory heptose metabolites. Therefore, we tested in a more targeted manner, whether the heptose gene cluster regulation and the output and activity of heptose reaction products would be modulated by mutation of the *cag*PAI (complete deletion mutants, leading to CagT4SS deficiency) (Fig. 3). We found that the complete isogenic deletion of the *cag*PAI had a significant, reducing effect on the transcript amounts, in particular of the *hldE* and HP0859 (*hldD*) genes, and partially on other genes of the cluster (qPCR, Fig. 3A to 3C). This observation was independently verified to be similar in three strains (26695a, Su2 and L7). The same pattern of downregulation of heptose biosynthesis transcripts was seen in transcriptome data of *cag*PAI deletion mutant in comparison to the parental wild type (Fig. 3D).

**Fig 3.**
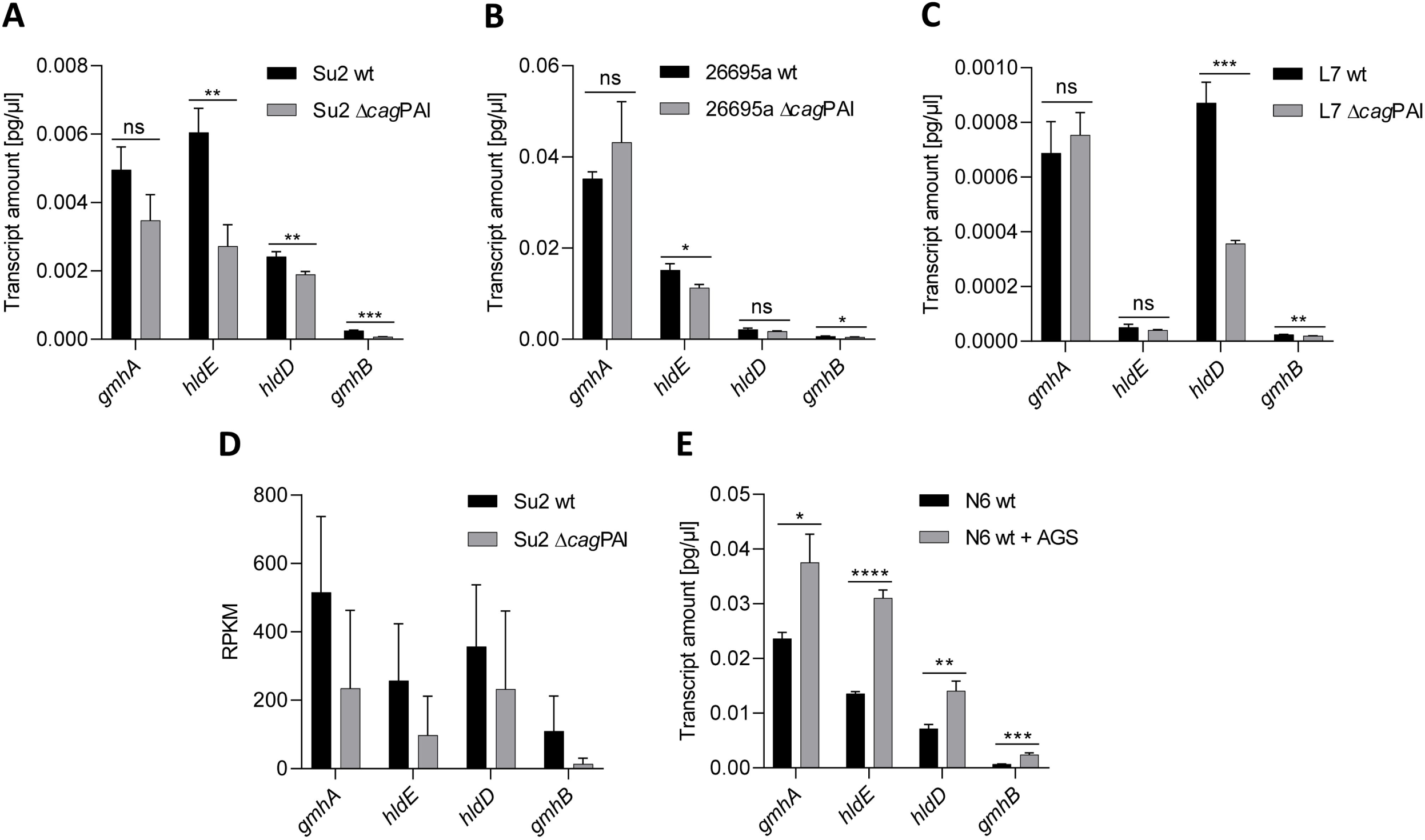
Regulation of heptose pathway transcripts in *H. pylori* is influenced by the presence of the *cag*PAI and contact with human gastric epithelial cells. **A)** Transcript quantification (RT-qPCR) of heptose biosynthesis genes of *H. pylori* Su2 wild type strain in comparison with its isogenic Δ*cagPAI* mutant. **B)** Transcript quantification (qPCR) of heptose biosynthesis genes HP0857 (*gmhA*) through HP0860 (*gmhB*) of *H. pylori* 26695a wild type strain in comparison with its isogenic Δ*cagPAI* mutant. **C)** Transcript quantification (qPCR) of heptose biosynthesis genes of *H. pylori* L7 wild type strain in comparison with its isogenic Δ*cagPAI* mutant. **D)** RPKM for genes *gmhA*, *hldE*, *gmhB* and *hldD* extracted from comprehensive transcriptome data for Su2 wild type strain or the Su2 Δ*cagPAI* mutant. **E)** Transcript amounts (qPCR) for genes *gmhA*, *hldE*, *hldD*, and *gmhB* of the heptose biosynthesis cluster (Fig. S1) of *H. pylori* N6 wild type co-incubated in presence or absence of AGS cells (MOI = 50) in cell culture medium for 4 h. All qPCR reactions were performed in triplicates and normalized against 16S rRNA transcript for each sample. qPCR results in panels A), B), C), E) are shown as absolute values (pg/µl), using 16S rRNA amounts and the respective wt samples as a reference for normalization. Statistically significant differences between conditions were calculated using student‘s *t*-test. Significances are marked with asterisks: * p < 0.05; ** p < 0.01; **** p < 0.0001; ns = non-significant.

Furthermore, we investigated how bacteria exposed to AGS cells in the presence or absence of a functional CagT4SS, regulated and expressed the LPS heptose biosynthesis gene cluster, using qPCR (Fig. 3E). Interestingly, we quantitated a significant upregulation of all genes of the heptose gene cluster in wild type and *cagPAI-*deficient bacteria that were associated for 4 h with AGS cells. At 8 h of co-incubation, the wild type bacteria maintained a stronger upregulation (Fig. S3) than the CagT4SS-deficient mutant.

### Transcriptome analyses of *H. pylori* in the presence or absence of the *cag*PAI reveal numerous transcript changes partially associated with carbon starvation regulator CsrA

Bacterial transcriptome changes have the potential to provide a comprehensive perspective on gene expression. Therefore, we performed whole bacterial transcriptome analyses on *H. pylori*, comparing the parental strain with an isogenic *cag*PAI deletion mutant, which we hypothesized to mimic a state of intrabacterial ADP-heptose enrichment by absence of the CagT4SS. Conditions of a “closed” or non-functional T4SS, which we had shown to reduce the expression of heptose genes in three different strains (Fig. 3), can also be helpful in identifying regulators or regulons possibly involved in a feedback regulation on heptose biosynthesis. The transcriptome analyses indeed revealed an influence of *cag*PAI deletion on numerous genes of different functional categories (Fig. 4A; Table 5, Suppl. Table S2). Using a cut-off of 1.5-fold regulated, about 500 genes were found both up- or down-regulated (Suppl. Table S2). Those included genes of different KEGG categories and also genes of the heptose biosynthesis gene cluster, with HP0858/*hldE* and HP0860/*gmhB* genes being significantly downregulated in single differential expression analyses (information on complete regulation dataset, see methods and Suppl. Table S2). Upon comparison with known regulons, a number of previously reported CsrA-dependent transcripts in *H. pylori* (compare to gene table in (46)), including transcripts of motility- and metabolism-associated genes, were differentially regulated under those conditions (Fig. 4A). CsrA broadly shifts metabolic, motility and other functions in responses to environmental conditions in various bacterial species (46–48, 53). Therefore, we also quantitated in detail the selected CsrA-related transcripts, *csrA*, fructose-bisphosphatase (*fbp*), riboflavin synthase subunit (*ribF*), and *acpP* using qPCR, comparatively between wt and *cag*PAI mutant, and confirmed the significant regulation of most of the transcripts (Fig. 4B) between both conditions. We then also compared selected, differentially regulated, CsrA-dependent transcripts with incubation of bacteria in contact with AGS cells, which impacts on the heptose biosynthesis gene cluster transcripts, as shown above (Fig. 3E), using qPCR. Thereby, we found indeed *acpP*, *csrA* and the CsrA downstream genes *fbp* and *rpoN* significantly up-regulated in bacteria co-incubated with gastric epithelial cells for 4 h (Fig. 4C). Generating a *csrA* mutant, we also verified that the heptose cluster and selected CsrA regulon transcripts are indeed dependent on CsrA function, by confirming their differential expression between parental strain and *csrA* mutant by qPCR (Fig. 4D, Fig. 4E). As a correlate of heptose biosynthesis with relation to pro-inflammatory cell activation, we finally demonstrated that *csrA* mutants show significantly lower activities than parental wild type bacteria on NF-κB luciferase reporter cells (Fig. 4F), suggesting that not only transcript amounts but also cell-active heptose metabolite MAMP production are reduced in the mutant. Taken together, the findings demonstrated that absence of the T4SS or human cell contact exerted an inversely correlated influence on the bacteria including regulation of heptose transcripts and other functions, in a CsrA-related manner.

**Fig 4:**
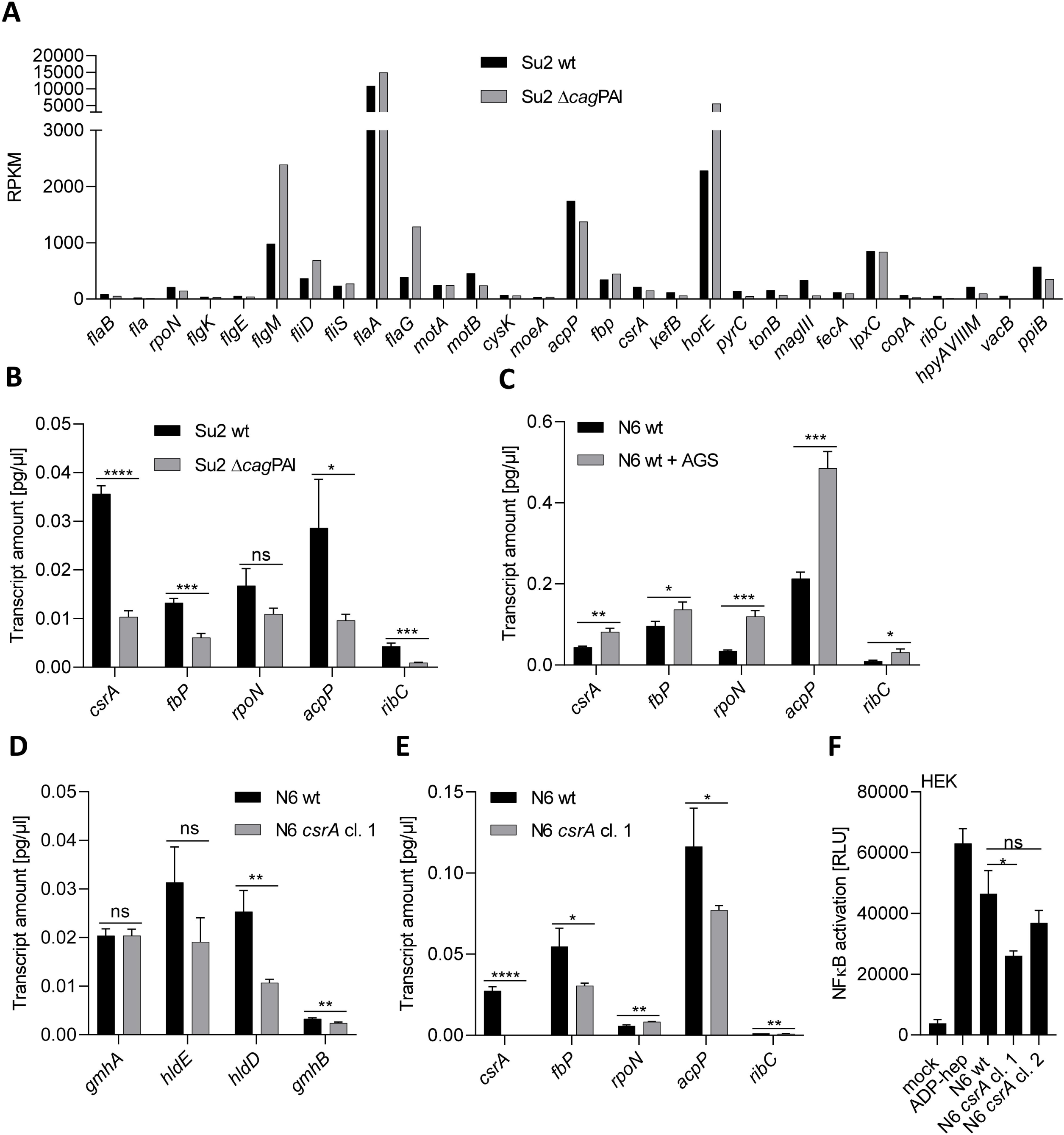
Heptose biosynthesis genes and other functional gene categories in the absence of a functional CagT4SS or in the presence of cells are regulated in association with the carbon starvation regulatory protein CsrA and in *csrA* mutants. **A)** Selected RNA-Seq results depicted as RPKM values from a transcriptome analyses comparing *H. pylori* wild type (wt), strain Su2, and its isogenic *cag*PAI deletion mutant. A representative experiment (R1) of two independently performed biological replicates is shown. The comparative data of the two replicates is fully listed in supplemental Table S2. Functionally annotated transcripts which are included in the CsrA regulon as defined in (46) are shown, representing functional categories of motility-related or metabolic genes. **B)** The same strains cultured under the same conditions were also compared for transcript amounts of CsrA-dependent genes (*csrA*, *fbP*, *rpoN*, *acpP*, *ribC*) by qRT-PCR. **C)** *H. pylori* (strain N6) cultured in the presence or absence of AGS cells were also tested by qRT-PCR for selected CsrA-dependent transcripts as defined under B). **D)** and **E)** A *H. pylori* N6 *csrA* mutant (allelic exchange insertion) was generated and specific gene transcripts of the heptose gene cluster and selected transcripts of the *H. pylori* CsrA regulon as in B) and C) were tested in qPCR. **F)** Reporter assays using HEK_NF-kB luciferase reporter cells (HEK) to detect a changed pro-inflammatory activity of *H. pylori* (strain N6) *csrA* mutants on the heptose dependent proinflammatory response. Two csrA mutant clones, cl. 1 and cl. 2, were tested alongside the N6 wt and reference pure ADP-heptose. Statistically significant differences between conditions were tested by unpaired student’s *t*-test. Significances are marked with asterisks: * p < 0.05; ** p < 0.01; *** p < 0.001; ns is non-significant.

**Table 5:** Part of comprehensive differential expression analysis for transcriptomes of *H. pylori* wild type (strain Su2) in the presence or absence of the *H. pylori* Δ*cag*PAI. The subset of the differential expression analysis show CsrA-dependent genes according to (46); only > 1.5-fold differentially regulated transcripts (ratio of Su2 Δ*cag*PAI vs Su2 wt [reference sample]) are included; a threshold of Max group mean of RPKM > 10 was applied. Gene name abbreviations are given in brackets; gene numbers are those of strain 26695 (39). Complete differential expression analyses (samples B13 and B14) are contained in Table S2.

| Gene name | Max group mean | Log<sub>2</sub> fold change | Fold change | P-value |
| --- | --- | --- | --- | --- |
| HP0172 (*moeA*) | 34.88 | 0.79 | 1.73 | 0.04 |
| HP0472 (*horE*) | 5,524.81 | 1.96 | 3.88 | 0 |
| HP0579 | 52.42 | -2.4 | -5.27 | 6.50E-03 |
| HP0580 | 67 | -1.39 | -2.62 | 1.42E-03 |
| HP0581 (*pyrC*) | 141.52 | -0.91 | -1.89 | 1.01E-03 |
| HP0601 (*flaA*) | 14,938.62 | 1.14 | 2.2 | 0 |
| HP0602 (*magIII*) | 333.82 | -1.77 | -3.42 | 4.91E-10 |
| HP0603 | 80.49 | -1.86 | -3.64 | 2.29E-03 |
| HP0629 | 54.18 | 0.95 | 1.94 | 8.87E-05 |
| HP0715 | 556.56 | 1.71 | 3.27 | 0 |
| HP0751 (*flaG*) | 1,284.04 | 2.4 | 5.29 | 0 |
| HP0752 (*fliD*) | 688.1 | 1.59 | 3.01 | 0 |
| HP0753 (*fliS*) | 270.58 | 0.87 | 1.82 | 4.53E-04 |
| HP0754 | 125.88 | -1.07 | -2.1 | 0.08 |
| HP0817 | 177.73 | -1.08 | -2.12 | 6.33E-03 |
| HP0906 | 38.85 | -1.78 | -3.43 | 6.72E-04 |
| HP1051 | 912.63 | -1.88 | -3.68 | 7.77E-16 |
| HP1052 (*lpxC*) | 848.04 | 0.67 | 1.59 | 1.58E-06 |
| HP1076 | 71.56 | -0.67 | -1.59 | 0.18 |
| HP1087 (*ribC*) | 53.09 | -1.32 | -2.5 | 0.01 |
| HP1122 (*flgM*) | 2,389.80 | 1.96 | 3.89 | 0 |
| HP1233 | 62.97 | -1.02 | -2.03 | 0.09 |
| HP1248 (*vacB*) | 56.09 | -2.02 | -4.04 | 1.18E-06 |
| HP1396 | 164.94 | 1.02 | 2.03 | 4.77E-06 |
| HP1439 | 13.65 | -0.81 | -1.75 | 0.61 |
| HP1566 | 57.96 | -3.7 | -12.97 | 8.50E-03 |

### Reconstitution of the *H. pylori* heptose biosynthesis pathway in vitro reveals various cell-active pro-inflammatory reaction products

The results of our regulation analyses and the observed strain differences in heptose cluster transcripts and HldE sequence and expression levels, prompted us to investigate which ones and which quantities of heptose products can be synthesized by the bacteria, resulting from the different reaction steps of the heptose pathway. In addition, transfection of *H. pylori* ETLs into reporter cells versus the medium co-incubation of reporter cells with ETLs revealed different responses, suggesting that more than one heptose product is present (see below). Using mass spectrometry, Pfannkuch and colleagues had so far found predominantly β-D-ADP-heptose (with trace amounts of β-D-heptose-1,7-bisphosphate - β-D-HBP) in treated lysates of *H. pylori* (one strain tested) (8). ADP-heptose is a cell-permeable metabolite (4, 8). β-D-HBP was suggested to be a second cell-active heptose metabolite, possibly produced in a strain-variable manner, which is not cell permeable (8, 41). Until now, other intermediate heptose metabolites of the pathway had not been identified in *H. pylori* or any other bacteria. To approach the question of various products of the *H. pylori* enzymes, we decided to reconstitute the reaction pathway of *H. pylori* in vitro, based on the three central, natively purified, *H. pylori* enzymes of the pathway (GmhA, HldE, GmhB; Fig. 5, Suppl. Fig. S4) from heterologous expression in *E. coli*. We also expressed and purified both functional domains of the bi-functional HldE enzyme (d1 and d2 domains; see Suppl. Fig. S1A, Fig. S4) separately for testing. Since the *H. pylori* HldE enzyme, due to its clear sequence heterogeneity between strains, was the eminent candidate for pathway and product heterogeneity within the biosynthesis pathway, we also expressed and purified the full-length enzyme from two different *H. pylori* strains (Fig. 5). We used these proteins in various one-pot in vitro reactions to convert seduheptulose-7-phosphate substrate in the presence or absence of ATP (see Methods). As an immediate read-out, we first tested the reaction products of different enzyme compositions on NF-κB luciferase reporter cells (HEK_luc epithelial cells) for determining cell-directed pro-inflammatory innate activation (Fig. 5A; Suppl. Fig. 4). Single purified enzymes, incubated without ATP and substrate, and heat-inactivated enzymes, were also tested in separate reaction mixes as controls, to exclude potential cell-activating contaminations after purification from *E. coli* (which all gave negative results; Suppl. Figure S4A).

**Fig 5.**
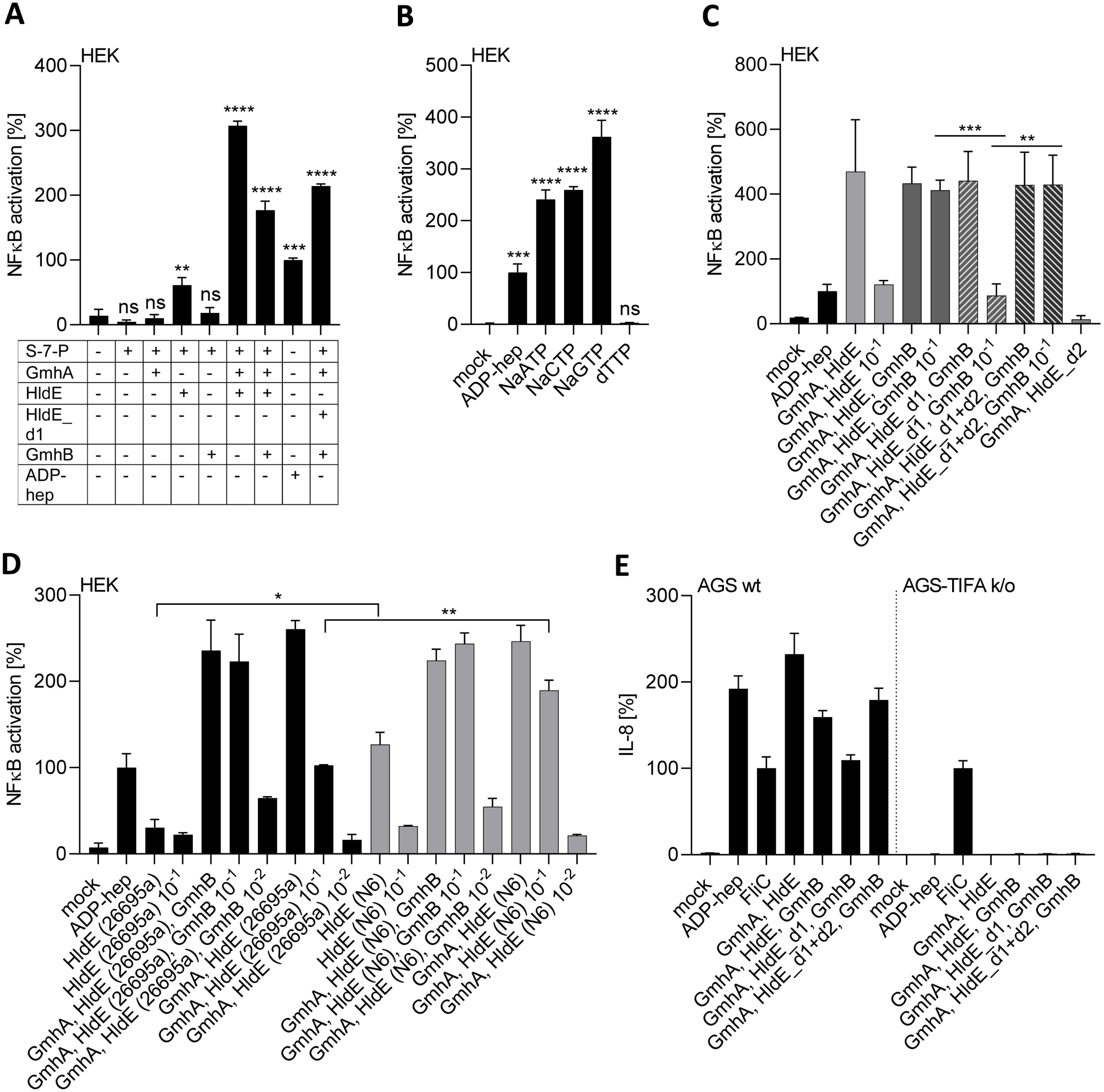
Reconstitution of *H. pylori* ADP-heptose biosynthetic pathway - *in vitro*-synthesized heptose metabolites of distinct enzyme combinations activate epithelial cells to a different extent. **A)** Pro-inflammatory activation of NF-κB reporter cells (HEK-NF-κB_luc) by heptose metabolites synthesized *in vitro* using recombinantly purified *H. pylori* proteins (strain 26695a) from substrate seduheptulose 7-phosphate (S-7-P). Absolute luciferase values are shown, and depicted in comparison to pure ADP-heptose (2.5 µM) co-incubated cells (control condition). **B)** HEK reporter cell NF-κB activation (normalized to 2.5 µM ADP-heptose condition, which was set to 100%) by products *in vitro*-synthesized using purified enzymes GmhA, HldE, GmhB (all from strain 26695a) and different nucleotides (all at 5.8 mM). **C)** HEK_luc reporter cell NF-κB-activation (normalized to ADP-heptose control condition, set to 100%) of *in vitro* reaction products synthesized by a combination of GmhA, GmhB plus HldE (latter as complete protein or as combined, separately purified HldE d1 and d2 domains, HldE_d1+d2), or HldE domain one (HldE_d1, from strain 26695a), or HldE domain two (HldE_d2) purified separately (non-diluted and 10^-1^-diluted reaction products are shown for each condition). **D)** NF-κB activation by products synthesized *in vitro* using the three-enzyme combination with highly purified HldE (complete protein) from two different *H. pylori* strains, 26695a or N6 as indicated on the y-axis (GmhA, GmhB remain in all reactions of strain 26695a). Strongly active mixes were added non-diluted, at 10^-1^, or at 10^-2^ dilution. The reporter cell assays in A), B), C) and D) were summarized from three biological replicates. **E)** Pro-inflammatory activation of AGS wild type (wt) and AGS-TIFA k/o cells by in vitro-synthesized heptose products, quantitated using IL-8 ELISA on cell supernatants. Values are shown in percent and were normalized to activation level of *Salmonella* FliC activation control (TLR5 ligand, 400 ng/well, set to 100%). The results shown in E) were summarized from two biological replicates for each experimental condition, each quantitated in technical triplicates. Statistical evaluation of significant differences was performed using unpaired student‘s *t*-test and shows statistically significant differences. Non-significant differences are not indicated. * p< 0.05; ** p < 0.01; *** p < 0.001.

Reaction products of the successful one-pot reconstitution with three main pathway enzymes combined (GmhA, HldE, GmhB) avidly activated cells (Fig. 5, Suppl. Fig. S4, Table 6). The concentration dependency of reaction products obtained *in vitro* was also established through titration in comparison with pure β-D-ADP-heptose (Suppl. Fig. S4B). We also reconstituted a reaction with only two purified enzymes, GmhA and HldE, expecting to obtain mainly β-D-HBP as reaction product. The reaction product(s) from this second reaction unexpectedly also activated cells strongly without the need for cell transfection (Fig. 5A, 5C). However, titrations showed that this two-enzyme mix was about one log less active than the three-enzyme mix (Fig. 5C). When incubated with other nucleotides than ATP, we obtained pro-inflammatory product with all nucleotides except dTTP in the three-enzyme mixes (Fig. 5B).

For HldE enzymes purified recombinantly from two different strains, which show considerable amino acid sequence diversity (Fig. S1B), both produced cell-active pro-inflammatory compounds. However, the enzymatic action of HldE from N6 resulted in a significantly stronger cell-activity in the two-enzyme reaction than HldE from strain 26695 (Fig. 5D). Even for HldE-only reaction mixes, we were able to produce cell-active output (Fig. 5D). In this case, the resulting activity was almost nil for 26695 HldE and also significantly higher for HldE from N6 strain (Fig. 5D). For all reaction mixes, we verified whether the resulting activity is dependent on the ALPK1-TIFA pathway, by comparing their activity on AGS wild type and AGS-TIFA k/o cells (11) using IL-8 release (ELISA) as a read-out. The TIFA k/o cells showed no pro-inflammatory activation by our mixes, confirming lack of product activity in the absence of TIFA-mediated signaling, while an alternative innate NF-κB stimulant, *Salmonella* FliC flagellin (TLR5 ligand), was positive (Fig. 5E).

**Table 6:**
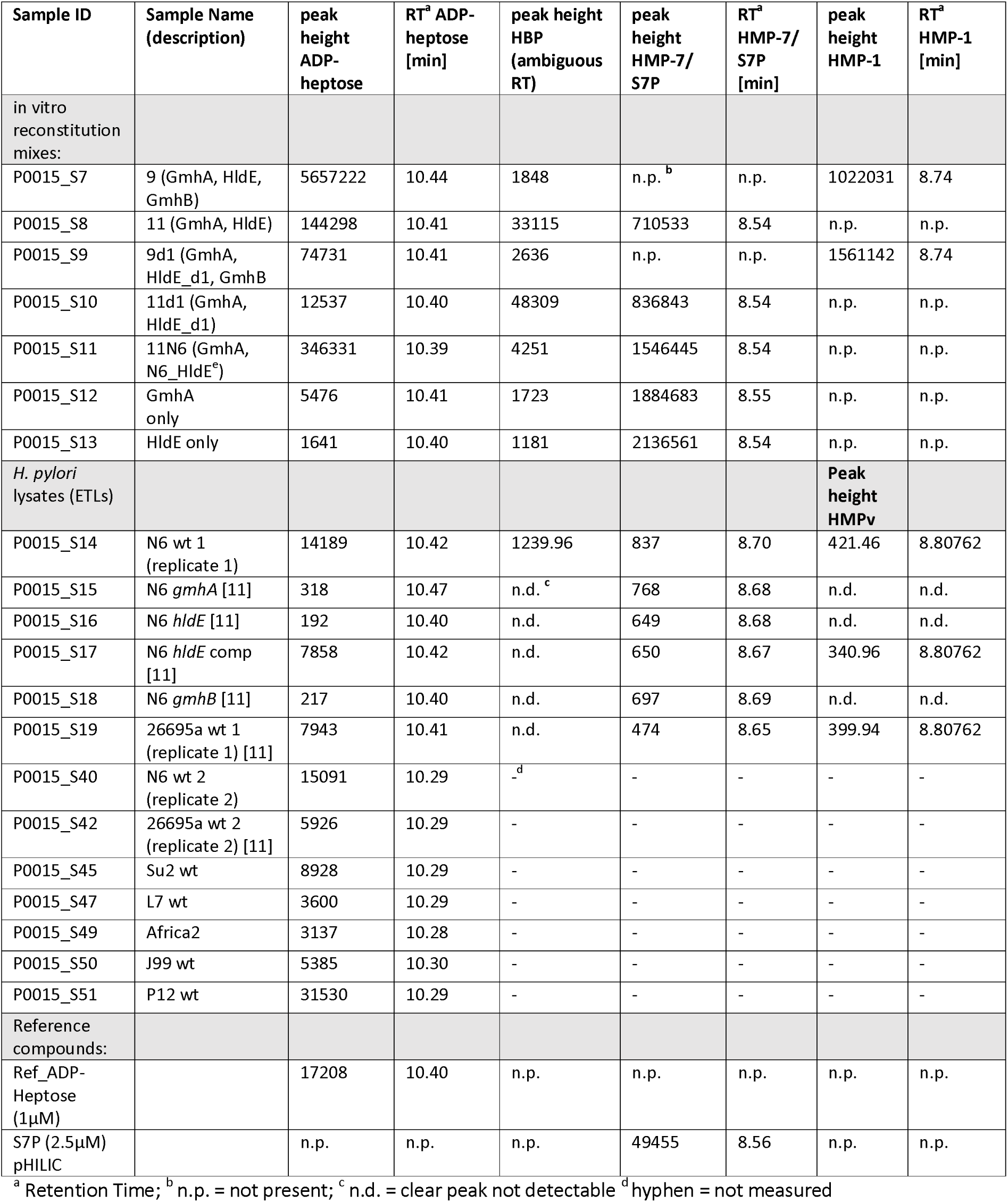
Sample identifiers for NMR and mass spectrometry analyses, and identification and quantification (chromatogram peak heights) of heptose metabolites detected by LC ESI MS-MS in in vitro reconstitution mixes and bacterial lysate (ETL) samples. In vitro reconstitution mixes (methods) contain purified enzymes from H. pylori strain 26695a if not indicated otherwise.

### Pro-inflammatory heptose pathway products identified as β-D-ADP-heptose and β-HMP-1 using NMR and mass spectrometry

To assign compound structures to the one-pot reaction products, we then used ^1^H NMR analysis of the reaction mixes containing different sets of recombinant enzymes from *H. pylori* (Fig. 6). Based on the specific chemical shifts, coupling constants and scalar correlations observed in two-dimensional COSY experiments, β-D-ADP-heptose was identified as the main reaction product of the combined action of GmhA, HldE, and GmhB proteins, when ATP was supplied as the energizing nucleotide (Fig. 6G). Comparing with reference compounds (β-D-ADP-heptose and β-L-ADP-heptose; confirmed by NMR, see Fig. 6E, Fig. 6F) we showed that the ADP-heptose product of the *H. pylori* enzyme reconstruction reaction was of the D-form (ADP-**D**-glycero-ß-D-manno-heptose) (Fig. 6G). In the two-enzyme reaction (GmhA and HldE), NMR spectrometry identified another, unexpected, cell-activating product, which was not identical to β-D-HBP but identified as predominantly β-HMP-1, in addition to the reaction mix ingredients (Fig. 6H). Dilution titration of the two-enzyme reaction mix containing β-HMP-1 yielded about ten-fold lower activity on cells (without transfection) as compared to the three-enzyme reconstitution (Supplemental Fig. S4B). Since previous studies had suggested that β-D-HBP is preferentially produced by HldE, we supplemented the HldE d1 and d2 domains (Fig. S1A, Fig S4) separately in the reconstitution. D2 domain alone, in combination with GmhA only (Fig. 5C), or with both GmhA and GmhB (not shown) did not yield any active product. Using solely the d1 domain of HldE enzyme, in combination with GmhA, we obtained a pro-inflammatory product that we also identified as β-HMP-1 by NMR, and no detectable β-D-HBP, suggesting that the HldE d1 domain activity was sufficient for HMP-1 production (Fig. 5C, Fig. 6I). D1 and d2 together behaved similarly as the complete HldE enzyme in all reaction mixes, yielding mainly ADP-heptose and a slight residual amount of β-HMP-1, but no detectable β-D HBP (Fig. 6J). The alternative nucleotides Na-GTP and Na-CTP, supplemented instead of ATP in the three-enzyme mixes, did not give detectable amounts of nucleotidyl-heptose despite pronounced pro-inflammatory cell activity; instead, β-HMP-1 was also revealed by NMR (not shown).

**Fig 6.**
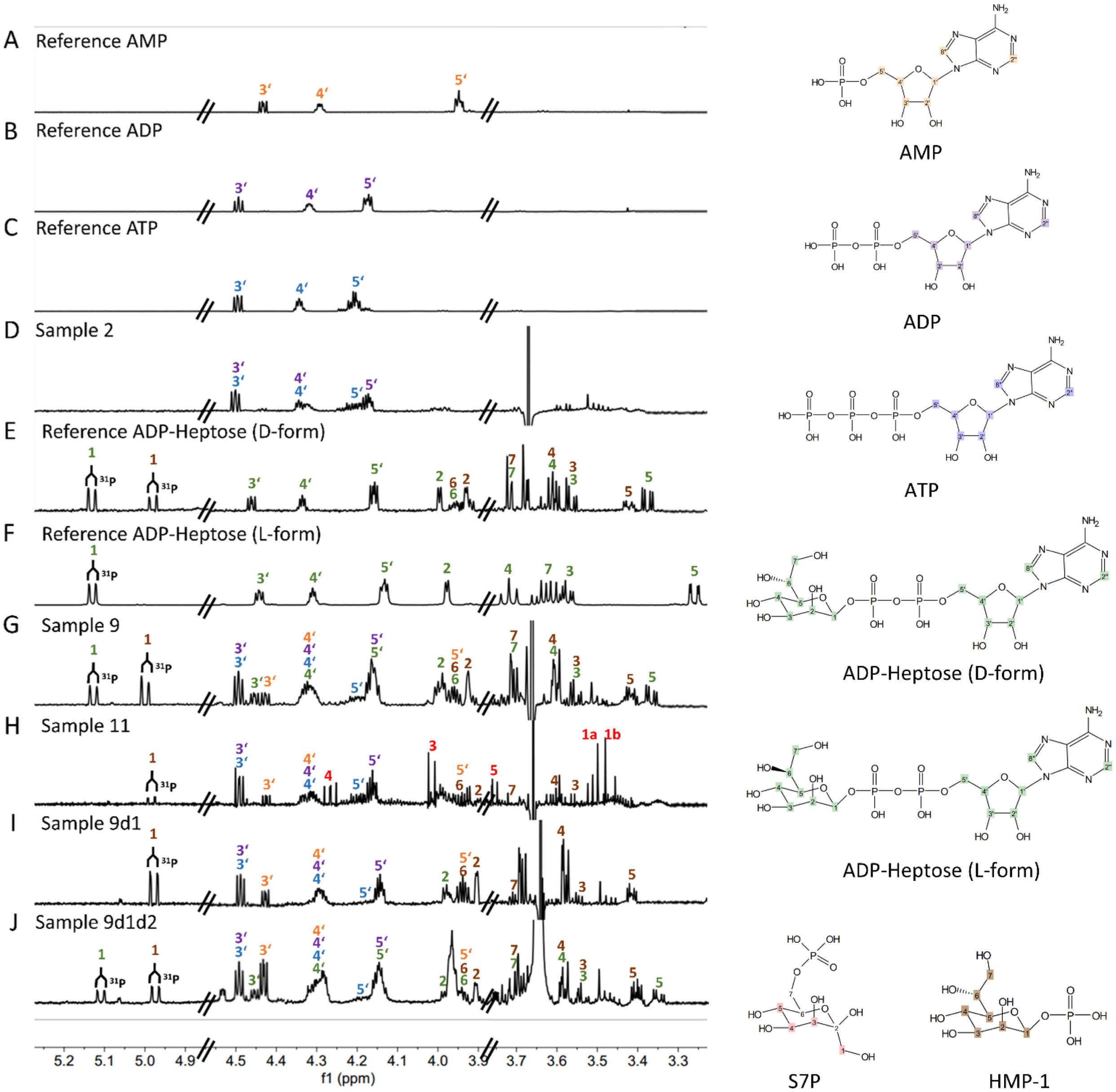
NMR analysis of reaction products of in vitro reconstituted *H. pylori* ADP-heptose biosynthesis pathway identify new cell-active product β-HMP-1. In vitro reconstitution of *H. pylori* ADP-heptose biosynthetic pathway was performed in one-pot reaction mixtures with various enzyme combinations as described in Fig. 5, Supplemental Fig. S4, Table 6 (sample identities), and in the methods. NMR was performed as described fully in the methods section. Panels **A)** to **F)** show the control conditions and control compounds as indicated in the upper left corner of each panel, analyzed in pure form (ATP, AMP, ADP, β-D-ADP-heptose, β-L-ADP-heptose; sample 2 (D) is a control mix without any substrate) Nomenclature of one-pot reactions (panels **D)**, **G)** to **J)**) was as follows: sample 2 - control sample with ATP and three enzymes, but no substrate; sample 9 – three enzymes, GmhA, HldE, GmhB, ATP and sedoheptulose-7-P (substrate); sample 11 - two enzymes, GmhA, HldE, ATP and substrate; sample 9d1 - three enzymes with domain 1 of HldE only, ATP and substrate; sample 9d1d2 – three enzymes with HldE d1 and d2, purified and added separately, ATP and substrate. All enzymes used here were cloned from strain 26695a. Structures of compounds and reaction products and their respective atoms are indicated to the right of each panel; atoms are numbered and labelled in different colors. Color-coded numbered peaks in the histograms correspond to the respective atoms of the input and output compounds, shown on the right, that were detected.

Due to the intrinsically low sensitivity of NMR spectroscopy in conjunction with severe signal overlapping in the NMR spectra of crude extraction mixtures, we were not able to detect any heptose compounds in *H. pylori* lysates by NMR. We therefore turned to mass spectrometry (LC ESI MS-MS) to test both, *in-vitro* reconstitution mixes and *H. pylori* ETLs (sample overview in Table 6), to verify compound structures and refine their detection and relative quantitation. In the *in-vitro* reconstitution samples, depending on their composition, we were able to clearly detect and quantitate ADP-heptose (Fig. 7A), HBP (Fig. 7C) and the heptose-monophosphates (Fig. 7B). In these samples, the three heptose-monophosphates, HMP-7, S7-P, and HMP-1, can be clearly distinguished by different retention times (Table 6). Highest amount of ADP-heptose product was quantitated in the three-enzyme mix containing full-length HldE. Most HMP-1 was detected and quantitated in the three-enzyme mixes containing either full-length HldE, or HldE d1 domain only (Fig. 7B, Table 6). HBP was detected predominantly in two-enzyme mixes containing GmhA and HldE, or GmhA and HldE-d1 (Fig. 7C, Table 6).

**Fig 7.**
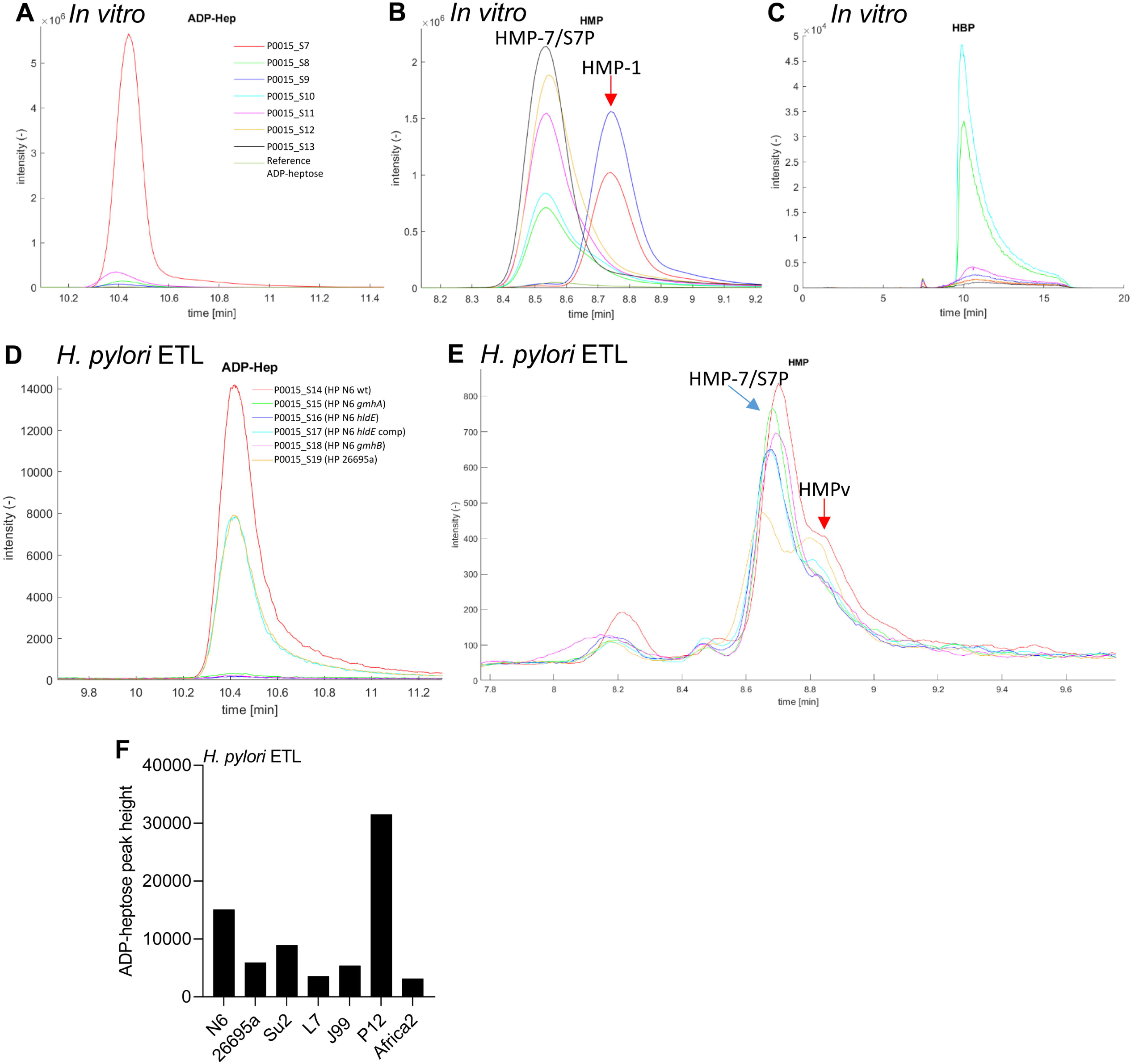
Mass spectrometry analysis (LC-ESI-MS-MS) confirms reaction products of *in vitro*-reconstituted *H. pylori* ADP-heptose biosynthesis pathway and identify new pro-inflammatory product β-HMP-1, and strain-specific quantities of ADP-heptose and other heptose metabolites in *H. pylori* lysates. In vitro reconstitution of *H. pylori* ADP-heptose biosynthetic pathway was performed in one-pot reaction mixes with various enzyme combinations (enzymes cloned from 26695a) as described in Fig. 5, Supplemental Fig. S4, Table 6 and in the methods. Mass Spectrometry was performed as described in the methods section. Panels **A)** to **C)** show the detection and quantitation of ADP-heptose (**A**), HMP variants (**B**) and HBP (**C**) in various one-pot mixes of purified *H. pylori* enzymes (see Table 6). **D**) and **E**) show the detection and quantitation of ADP-heptose and HMP variants by mass spectrometry in *H. pylori* lysates (ETLs). Same color-codes of chromatogram curves in the panels correspond to the same samples in A), B), C), respectively, and in D) and E), respectively. In D), samples S14 (N6 wt ETL), S17 (N6 *hldE* comp ETL) and S19 (26695a wt ETL) show strain-specific, high amounts of ADP-heptose. In E). The same samples show also an increased shoulder peak, in comparison to the other samples, at the approximate retention time of heptose-monophposphate-1 (HMP-1). This shoulder is designated here as heptose monophosphate variant, not identical to HMP-7 (HMPv), since the assignment of the shoulder peak was ambiguous. Peaks in the chromatograms are labelled with the respective compound designations (input and output compounds). See Table 6 for full descriptions of samples, compound identifications and quantifications (peak height).

Mass spectrometry also detected a distinct heptose monophosphate species (HMPv, probably HMP-1; Fig. 7E) as a novel compound (shoulder peak in chromatogram, matching the retention time in reconstitution mix containing HMP-1) in addition to ADP-heptose (Fig. 7D), both in *H. pylori* wild type lysates and ETL of an HP0858 (HldE) overexpression strain (Fig. 7E, Table 6). Interestingly, greatly divergent amounts of specific metabolites in two different wild type isolates were quantitated by LC-MS, with respect to amounts of ADP-heptose and the probable HMP-1 monophosphate species (Table 6). Differences in ADP-heptose quantities between N6 and 26695a strain lysates were more than 20-fold (Table 6), and ADP-heptose content in lysates also varied quantitatively for a broader range of diverse wild type clinical isolates (Fig. 7F). Similarly, proposed HMP-1 heptose-monophosphate showed quantitative differences between the two strains (Table 6). This clearly corresponded with a significant difference in pro-inflammatory cell activity between both strains (live and lysates; Fig. 2). HBP, while clearly revealed in enzymatic reconstitution mixes (Table 6), was poorly detectable by LC-MS in bacterial lysates, probably due to low concentrations and compound instability.

### Medium supplementation versus cell transfection of reaction products of in vitro reconstitution and *H. pylori* ETLs activate cells differently: further evidence for differentially active heptose products

When we co-incubated reporter cells with reaction products from the in vitro reconstitution reactions, we observed strong pro-inflammatory cell activation, using the three-enzyme reconstitution (above), mostly producing β-D-ADP-heptose according to NMR results. This activity was absent when TIFA k/o cells were used, which cannot mount an innate response towards heptose metabolites (Fig. 5E), clearly assigning the activity to the heptose product(s). The cell-directed pro-inflammatory activity was markedly lower when co-incubating reporter cells with reaction products from the two-enzyme reaction (enriched in β-HMP-1 and β-D HBP) by merely supplementing the medium. In order to gather more evidence for a mixture of active products in some of the reactions, we transfected pathway reconstitution products and bacterial ETLs into the HEK-NF-κB_luc reporter cell line (Fig. 8A). While transfection of the three-enzyme pathway product (containing mostly ADP-heptose as per our biochemical analyses) elicited about the same amplitude of response as co-incubating the same amount in the medium, the products of the two-enzyme reconstitution behaved differently; in this case, the transfection of the product elicited at least three-fold higher activity in the reporter cells in comparison to the medium-supplementation with the same reaction products (Fig. 8A). When analyzed using mass spectrometry, the two-enzyme reaction contained markedly higher amounts of the non-permeable metabolite β-D HBP than the three-enzyme mix (Fig. 8B, Table 6). We also compared the pro-inflammatory activity of bacterial ETLs collected from five different *H. pylori* strains (26695a, Su2, N6, L7, and Africa2) and isogenic Δ*cag*PAI or *cagY* mutants of three strains (Fig. 8C, 8D) on cells with and without lipofectamine transfection. In those experiments, the ETLs always activated significantly more when transfected (Fig. 8C, 8D). Strain differences were determined for the same amount of ETLs using the ratio between transfected and non-transfected, quantitating how strongly the activation was increased by transfection. The ratios ranged from slightly above or below two-fold increase for N6 and Su2 (Fig. 8E), to between three-fold and four-fold for 26695a, L7 and Africa2. This corresponds to our analyses of ETLs containing a) cell-permeable ADP-heptose, (shown before in (8)), b) an pro-inflammatory active monophosphate compound (HMPv), likely HMP-1, which also has a good cell permeation activity (41), and c) at least one other heptose metabolite, which is less cell-permeable (suggested to be HBP (5, 8)). In vitro one-pot reactions which cannot produce β-D-HBP (e.g. a mix including GmhA, d1 domain of HldE, and GmhB), and which contained HMP-1 (by NMR and LC-MS) as the only active product, did not result in an increased activation upon cell transfection. We also tested several isogenic heptose biosynthesis and Δ*cag*PAI mutant lysates in comparison with their wild type parent ETLs by cell transfection (three strains). Most mutants did not behave differently from their wild type parent (Fig. 8D, 8E). For strain N6, the isogenic *gmhB* mutant ETL (expected to increase HBP product) displayed a substantial activity increase by transfection over non-transfected (Fig. 8D), and a net increase of activity over the parent strain. Interestingly, testing the 26695a *cagPAI* deletion mutant ETL revealed a stronger NF-κB activity increase by transfection over non-transfected than the 26695a wild type ETL (Fig. 8E). *cagPAI* deletion would mimic a closed T4SS. Those results strongly suggested that *cagPAI* presence can affect the production of metabolites which are less permeable than ADP-heptose.

**Fig 8.**
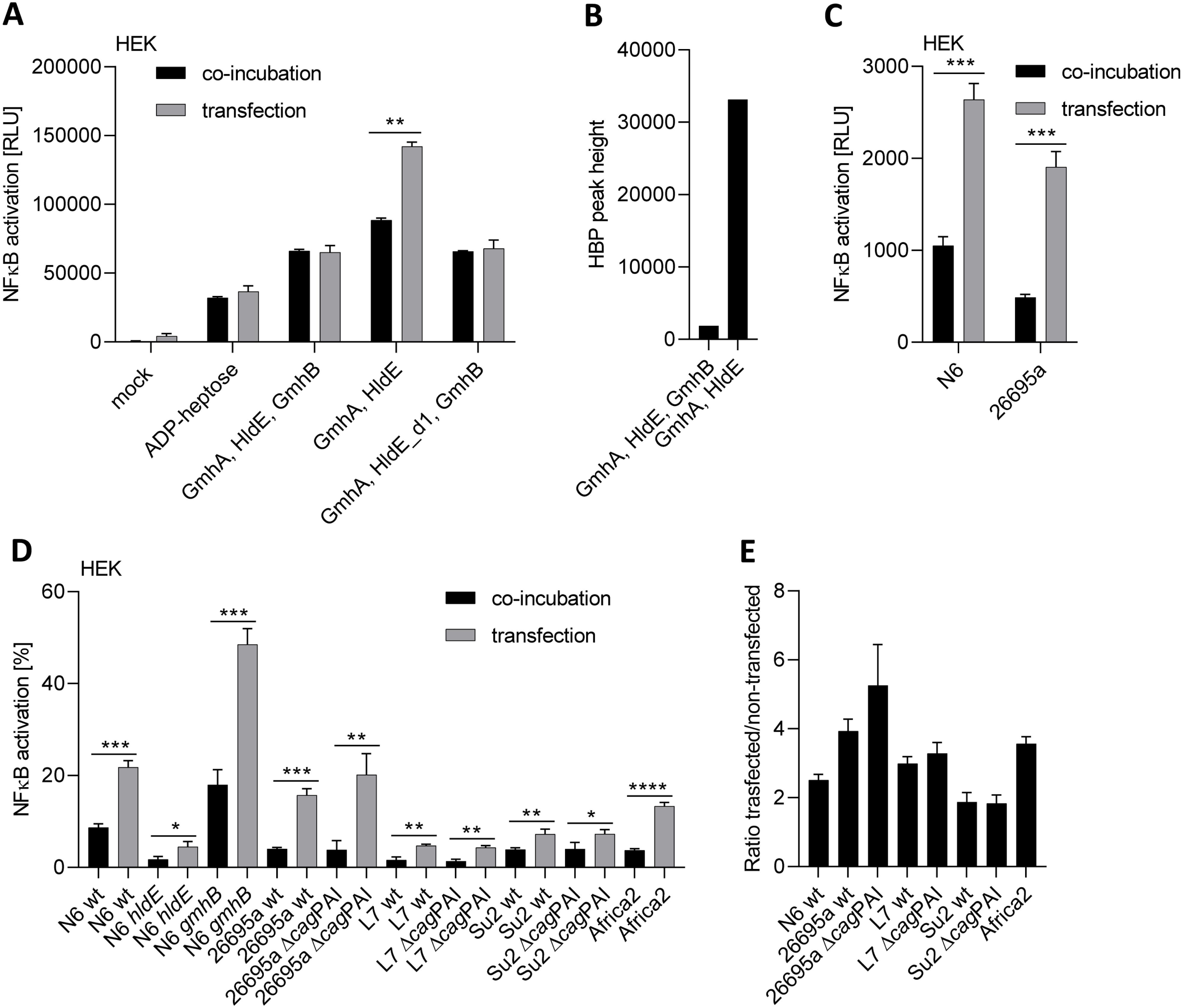
Transfection improves the pro-inflammatory activity of partially reconstituted heptose reaction (GmhA, HldE) and of some bacterial ETLs. **A)** Activation of HEK-NF-κB_luc reporter cells after co-incubation or transfection by reconstituted heptose mixes produced using the recombinant *H. pylori* enzymes GmhA, HldE (complete enzyme or separate domain d1) and GmhB in different combinations (4 h cell co-incubation). **B)** shows the comparative quantitation of HBP by LC-MS-MS mass spectrometry in the reconstitution mixes of three heptose pathway enzymes versus two enzymes (correspond to samples S7 and S8 in Table 6). The three-enzyme mix contains very little HBP, while the two-enzyme mix shows a more than 10-fold increase in HBP. **C)** Activation potential of *H. pylori* ETLs on HEK-NF-κB_luc reporter cells after co-incubation or transfection of enzymatically treated lysates (ETLs) of the *H. pylori* wild type strains N6 and 26695a (4 h co-incubation). Reporter activation is quantitated in absolute luminescence values (RLU) in A) and C). **D)** Differences in proinflammatory cell activation (HEK-NF-κB_luc) between co-incubation with and transfection (T) of various *H. pylori* strains’ ETLs. Strain and mutant designations (wild type = wt of strains N6, 26695a, L7, SU2 and Africa2, mutants designated by respective gene names) conform to previous figures. Incubation time was 4 h for all conditions; all values (shown in %) were normalized to ADP-heptose co-incubation of the same experiment (2.5 µM, not shown), which was set to 100%. Experiments to detect NF-κB activation in A), C) and D) were performed in to (A) or three biological replicates each and were repeated at least once on different days. Statistically significant differences between conditions of co-incubation and transfection were calculated using student‘s *t*-test. Significances are marked with asterisks: * p < 0.05; ** p < 0.01; *** p < 0.001; **** p < 0.0001. **E)** Calculation of ratios between co-incubated and transfected conditions for response to selected strains’ ETLs (only wt strains and isogenic T4SS-deficient mutants, same co-incubation conditions as in D)).

## Discussion

Heptose metabolites from diverse bacteria have recently been characterized as MAMPs/PAMPs with strong pro-inflammatory activating potential for human myeloid and epithelial cells (12, 49). Some heptose metabolites such as ADP-heptose can be actively taken up as solutes after they have been released into the environment (5, 7, 8, 41). A number of pathogenic bacteria including *H. pylori* and Enterobacterales do not release a substantial amount of heptose metabolites into the medium, but mainly use their host-directed secretion systems (T4SS, T3SS) for direct transfer of the metabolites into host cells (4, 8, 11, 12, 14). This mode of targeted metabolite transport raises the obvious question, whether and how the bacteria employ active mechanisms to regulate the activating metabolite biosynthesis, possibly in a strain- and cell contact-specific manner. Envisaged mechanisms would comprise specific sensing of environmental cues and subsequently lead to increased activation of heptose biosynthesis genes upon host cell contact.

Using the model organism *H. pylori*, which translocates ADP-heptose and possibly other heptose metabolites via its type 4 secretion system, we have therefore addressed several questions regarding strain diversity in heptose gene regulation, diverse heptose metabolite generation, and variable pro-inflammatory effects of bacterial heptoses on host cells. First of all, we obtained a fundamental knowledge about strain diversity in transcript amounts and transcriptional and post-transcriptional regulation mechanisms of the *H. pylori* heptose biosynthesis gene cluster. Transcript amounts varied between strains and between the genes of the cluster in each tested strain, with the last cluster genes, and most relevant genes for generating pro-inflammatory heptose intermediate products (11), *gmhA* and/or *hldE*, frequently possessing the highest transcript activities. Both, the absolute transcript amounts of each gene and relative expression patterns of the cluster genes were extraordinarily strain-variable. In particular, we found strains with very high transcript amounts specifically of *gmhA* and *hldE*, while other strains such as L7 (Asian origin) and Africa2, a primary *cag*PAI-negative strain, had higher transcript amounts selectively of both *gmhA* and *gmhB*, but not of *hldE*. Those observed expression differences can explain some of the strain differences in pro-inflammatory bacterial lysate activity and in cell responses induced by live bacteria that have been observed in this work and before (45). In addition, while *gmhA, gmhB* and *hldD* did not differ markedly in sequence between *H. pylori* strains, the bifunctional *hldE* gene, crucial for the generation of pro-inflammatory cell-active heptose metabolites (3, 10, 8, 11), showed a comparably strong inter-strain DNA and derived amino acid sequence variability. Interestingly, using a specific antiserum against HldE, we could also pin down marked strain-specific differences in HldE protein content. The finding that the different strains had a very diverse heptose-dependent pro-inflammatory activation potential on epithelial cells ((11, 45), and our new results), is an expected outcome of expression differences and HldE sequence diversity. HldE protein expression in different mutants (*cagY*, Δ*cag*PAI) of the same strain appeared not much changed, except for the HldE mutant. Taken together, our findings of transcript profile diversity and strain-variable HldE protein content suggest a major role of this bi-functional enzyme in host interaction. Since bacterial lysates do not permit to detect metabolite transport differences of live bacteria, we have still less insight into active metabolite production and content in live bacteria in the presence of target cells. Upregulation of the heptose cluster genes in wild type bacteria with an active T4SS in the presence of cells is highly suggestive of increased heptose metabolite biosynthesis upon cell contact.

In order to identify novel pro-inflammatory heptose metabolites that *H. pylori* can variably produce, and to identify strain-specific differences in enzyme activities, as one possible explanation for inter-strain variation of active metabolite content, we reconstituted the heptose biosynthesis pathway in one-pot reaction mixes. We used purified HldE enzyme from two different *H. pylori* strains (26685 or N6), and two separate HldE domains. When we tested the metabolic reaction products on NF-κB reporter cells, the fully reconstituted pathway mix (three enzymes, GmhA, HldE, and GmhB, combined) as well as the combination of GmhA and HldE (two enzymes only) both produced pro-inflammatory cell-active metabolites, although the latter reaction cannot form ADP-heptose. Transfection of reporter cells by pathway reconstitution products revealed that the two-enzyme reaction products acted much stronger upon transfection, while the three-enzyme product (predominantly β-D-ADP-heptose and HMP-1 according to NMR and mass spectrometry analysis) showed no difference between transfection and supplementation in the cell medium. The *H. pylori* pathway had been only partially enzymatically reconstituted *in vitro* prior to the present study, using synthetic β-HMP-7 or β-HMP-1 as substrates, and GmhB and/or HldE as purified enzymes (8). In the latter study, only β-D-ADP-heptose was identified as the reaction end product in *H. pylori* lysates (using mass spectrometry), which is cell permeable, while β-D HBP, an intermediate pathway product, is not (5). Reconstituted heptose pathway from *Campylobacter jejuni*, in a three-enzyme mix, was previously assessed to activate cells by external supplementation (7), while the latter study did not characterize the reaction products further. We established differences in the processive enzyme activities of HldE proteins from two different strains (26695a and N6), which show considerable amino acid sequence diversity, using the one-pot mixes, followed by LC-MS analysis of the products. HldE of strain N6, alternatively added to the enzyme mixes, was significantly more active, even as a single enzyme or in the two-enzyme combination with GmhA and produced significantly more ADP-heptose detectable by mass spectrometry, both in the enzyme mixes and in *H. pylori* lysates (Fig. 7D). The findings of inter-strain transcript, regulatory as well as enzyme variation in the heptose biosynthesis pathway points to a variety of factors which can influence the strain diversity in cell-directed metabolites and pro-inflammatory activity.

In the present study, we could now also unequivocally assign major reaction products formed in one-pot reaction mixes containing recombinant *H. pylori* enzymes, and in *H. pylori* lysates, using both NMR spectroscopy and mass spectrometry. In the fully reconstituted mix, we identified mainly β-D-ADP-heptose and HMP-1. In the two-enzyme mix, we identified β-D-HBP by mass spectrometry, and in addition, high amounts of a second, pro-inflammatory metabolite heptose-monophosphate variant, β-HMP-1, which had been synthesized in vitro as novel cell-active heptose product before (41) but not yet identified in bacteria directly. β-HMP-1 can theoretically be formed already in the second enzymatic step of the biosynthesis reaction, which we could confirm in a mix using only the HP0858 d1 domain in addition to GmhA. In *H. pylori* lysates, depending on strain and mutant identity, we quantitated as main cell-active products both ADP-heptose and a probable monophosphate (likely HMP-1), which was never before identified directly in bacteria. Particularly intriguing were the strong differences in amounts of ADP-heptose and pro-inflammatory heptose-monophosphate quantities between tested wild type strains. From transfection experiments of lysates and the two-enzyme preparations, we assume that, additionally, a third active compound other than ADP-heptose and probable HMP-1 is contained. HBP, which was only detectable in one tested strain (N6) by mass spectrometry, had been detected before in low quantities, and found not to be cell-permeable (4, 8, 41) We established that the activities and metabolite content of wild type lysates and *in vitro* reactions containing HldE proteins from two different *H. pylori* isolates (N6 and 26695), which show considerable amino acid sequence diversity, were different. HldE reaction mixes of strain N6 were significantly more enzymatically active, even as a single enzyme, or in the two-enzyme combination with GmhA, to produce HMP-1 or ADP-heptose. This matched to the higher detectable content of metabolites in strain N6 bacterial lysate, which was only exceeded by another frequently used strain, P12 (8). Strain differences in the production of β-D-ADP-heptose and other heptose metabolites by *H. pylori* can therefore be detected, and partially quantitated, directly in lysates. Strain differences in heptose metabolite production can also be inferred in our present study from a) distinct transcript amounts of various heptose cluster genes between different strains, b) varying enzyme activities, or c) the strain-variable ratios between lysate-transfected and co-incubated conditions. In any case, for the production of cell-active intermediate pathway metabolites, the central bi-functional enzyme HldE is essential. Through our in vitro reconstitution, we also collected evidence that domain 1 of HldE might, strain-specifically (e.g. in N6), not only act as a kinase but also as a phosphatase *in vitro*. This finding may resolve the conundrum that in distinct live *H. pylori* isolates, the phenotype of heptose metabolite-dependent cell activation of HP0860 (*gmhB*) mutants was found to be strain-dependently distinct (8, 11). Since we demonstrated that the *H. pylori* heptose biosynthesis pathway is able to produce a pro-inflammatory intermediate metabolite HMP-1 in the first two steps already with only GmhA and HldE, the next enzymatic steps may, in principle, not be necessary to produce pro-inflammatory metabolites.

Expressed separately, the two domains of the HldE enzyme, corresponding to the different enzymatic domains, had differential effects when used as part of the reconstitution reaction – single d2 addition in the three-enzyme mix did not lead to any cell-active product, while single d1 domain addition also produced detectable cell-active β-HMP-1 (NMR and mass spec). Activity of the latter reaction product was not higher when transfected into cells, different from the products of the GmhA and HldE two-enzyme reconstitution, supporting again the notion that a non-permeable product (HBP) causes the additional activation. The use of different nucleotides as cofactors in the reaction revealed that nucleotides are essential cofactors for all steps of the reactions to proceed. Without nucleotides, no products are revealed. Secondly, nucleotides other than ATP (CTP, GTP) can serve as cofactors of the enzymes in the first two reaction steps as efficiently as ATP, but cannot be transferred into nucleotidyl-heptose by HldE, as the reaction product then stopped at β-HMP-1. Adekoya et al. (41) demonstrated that, when provided at equal molarities with transfection, β-HMP-1 was about equally active on HEK epithelial cells than β-D-HBP, while HMP-7, the primary intermediate reaction product generated from S-7-P by GmhA, also detected in some of our samples, was not cell active at all. Our co-incubation results also confirmed that various human cell types are able to take up ADP-heptose and β-HMP-1 metabolites from the medium, without the need of a bacterial transport system ((12) and present study).

In addition to strain-specific traits in transcript amounts, enzyme and lysate activities of the heptose biosynthesis gene cluster, we identified a role of presence and activity of the *cag*PAI and the CagT4SS. In mutants not possessing the *cag*PAI, the heptose cluster genes were less expressed than in the corresponding wild type strains. A closed or absent secretion system, inducing feedback regulation, broadly influenced bacterial gene regulation, time- and strain-dependently, also leading to the downmodulation of heptose pathway transcripts. Presence of cells also modulated bacterial gene regulation and, in turn, upregulated heptose pathway transcripts. Our regulation analyses therefore suggest that heptose cluster transcript activities and pathway output can be adjusted according to T4SS activity and presence of cells. Prompted by the pathway analysis of comprehensive differential transcriptomes in *cag*PAI-deleted bacteria, we obtained evidence that the modulation of the heptose gene cluster in *H. pylori* is at least partially under CsrA regulation (46, 50–52). CsrA is an important metabolic and global posttrancriptional bacterial regulator on RNA, which influences broadly metabolic pathways and motility functions in various bacteria (50–53). A preliminary definition of a genome-wide set of CsrA-dependent transcripts in *H. pylori* was performed before (46), and our comprehensive dataset of T4SS-dependent regulation in the present study shows some overlap. Recent work in *E. coli* has impressively demonstrated how CsrA acts on a genome-wide scale, both as an activator and as a repressor (53). We generated and characterized *H. pylori csrA* mutants, which demonstrated that the heptose gene cluster, in addition to known CsrA-regulated transcripts, is under CsrA control, and that *csrA* mutants also exert lower proinflammatory activity on human cells. As one signature gene downstream of CsrA, whose activity is governed by fluxes in carbohydrate metabolism (54), we also confirmed *fbp* (coding for fructose-bisphosphatase). *fbp* gene product, fructose 1,6-bisphosphate, is an upstream modulator of central metabolism and glycolytic flux (glycolysis and TCA cycle activation versus gluconeogenesis) in bacteria (54). We found *fbp* transcript upregulated alongside *csrA* upon bacterial co-incubation with human gastric epithelial cells. The results are suggestive of a global regulatory switch of *H. pylori* phenotype between ecological conditions (e.g. planctonic versus cell-associated) under which cell-directed heptose transport and other functions are either shut off or switched on.

In conclusion, *H. pylori* produces a pro-inflammatory heptose-monophosphate MAMP, most likely β-HMP-1, in addition to β-D-ADP-heptose and HBP. In addition, heptose biosynthesis pathway activity as well as HldE expression are regulated strain-specifically and dependent on *Cag*T4SS activity, which entails a partially CsrA-dependent regulatory feedback on heptose biosynthesis and global functions. Hence, intracellular heptose metabolite amounts and an active CagT4SS are likely to influence bacterial central metabolism and induce cell contact-dependent heptose production “upon” demand. Consequently, *H. pylori*, an important model pathogen and paradigm of persistently host-associated bacteria, increases its toolbox of mechanisms to influence and fine-tune its production of cell-active, translocated, pro-inflammatory carbohydrate metabolites and thereby possibly combines innate immune activation of host cells with metabolic crosstalk.

## Acknowledgements

We are grateful to Bettina Sedlmaier-Erlenfeld for expert technical assistance. We gratefully acknowledge the contributions of Tatiana Murillo during an internship. We thank the Deutsche Forschungsgemeinschaft for funding the project in the framework of the Center Grant SFB 900, project B6 to CJ and in the framework of EI 384/16-1 to WE. MH, FM and LF acknowledge support by the intramural Max von Pettenkofer Institute, LMU Munich, graduate program “Infection Research on Human Pathogens@MvPI”. We thank Sebastian Suerbaum for helpful comments on the manuscript and continuous support. HL and JR acknowledge funding from the Cluster of Excellence EXC 2124 from the Deutsche Forschungsgemeinschaft.

## Supplemental Material

**Supplemental Figures S1 through S4, Legends, and Supplemental Tables S1 and S2**

**Fig S1. Genomic arrangement, protein diversity and strain-specific regulation in the *H. pylori* heptose biosynthesis gene cluster. A)** Organization of heptose biosynthesis operon genes HP0861 to HP0857/*gmhA* in *H. pylori* 26695a. *hldE* is highlightes in green color. Gene arrangement is conserved in other strains isolated worldwide. **B)** variation of HP0858/HldE protein sequence in diverse *H. pylori* strains (B8, J99, V225, 26695, B38, Cuz20, F32, OK310, SA7, HPN6) selected from different geographic origins and strain populations. Note the very conspicuous hinge region in HldE between the two domains d1 and d2, characterized by gaps in some strains. **C)** to **G)** depict quantification of transcript amounts of heptose biosynthesis cluster genes *gmhA*, *hldE*, *gmhB*, and *hldD* of *H. pylori* wild type strains N6 (C), 26695a (D), Su2 (E), L7 (F) and Africa2 (G) by qPCR, performed in technical triplicates. All qPCR results, given in absolute quantities of pg/ml, were normalized to 16S rRNA transcript amounts of each sample. Pairwise significance of differences (p values) in panels C) through G) was calculated by unpaired student‘s *t*-test. Significance values: * p < 0.05; ** p < 0.01; *** p < 0.001; **** p < 0.0001; ns = non-significant.

**Fig S2. Gastric epithelial cell line MKN28 (Asian ethnic origin) response to *H. pylori* ETLs. A)** to **D)** Activation of MKN28 gastric epithelial cells after co-incubation with ETLs produced from various *H. pylori* wild type strains (A) and mutants (B to D) as indicated on the x-axis, for 4 h. A quantitative read-out for pro-inflammatory response was obtained by performing IL-8 ELISA. Shown are the results from technical triplicates of biological duplicates. All experiments were repeated at least once on two different days. In B), C) and D), statistical significance was calculated for differences between wt strain and each mutant or complemented strain. Significance of differences (p values) was calculated by unpaired student‘s *t*-test. Significance values: ** p < 0.01; *** p < 0.001; ns = non-significant. In all cell activation experiments, co-incubation with pure ADP-heptose (2.5 µM, shown in A)) served as a reference for activation.

**Fig S3. Time-dependent regulation of heptose biosynthesis gene cluster in *H. pylori* N6 and its isogenic *cagY* mutant, co-incubated with gastric epithelial AGS cells. A)** and **B)** show transcript amounts (RT-qPCR) of heptose cluster genes *gmhA*, *hldE* (HP0858), *hldD*, *gmhB* (HP0860) of *H. pylori* N6 wild type and HP0527 (*cagY*) mutant, both co-incubated in presence or absence of AGS cells (MOI=50) for 4 h (A) or 8 h (B), respectively. Control bacteria were incubated in cell culture medium alone for the respective time periods. Three technical replicates are summarized in the panels. All qPCR results were normalized to 16S rRNA transcript. Statistically significant differences (p) between conditions were calculated by unpaired student‘s *t*-test. Significance values: * p < 0.05; ** p < 0.01; **** p < 0.0001; ns = non-significant.

**Fig S4. Quality controls of purified recombinant *H. pylori* heptose biosynthesis enzymes and activity titrations for *in vitro* reconstitution of heptose biosynthesis pathway. A)** Activation of NF-κB reporter cells (HEK_luc) by heptose metabolites synthesized using one-pot reactions of recombinant *H. pylori* 26695a proteins with enzymatic substrate seduheptulose 7-phosphate (S-7-P), and respective controls. Controls include active single enzymes plus substrate, or heat-inactivated (HI) enzymes plus substrate. **B)** Activation of HEK NF-κB luciferase reporter cells by pure β-D-ADP-heptose (titration), the products of three-enzyme combination GmhA, HldE and GmhB, the reaction product of two-enzyme combination GmhA, HldE (in vitro reconstitution: in each case non-diluted reaction sample and the same used at different dilutions from to 10^-1^ to 5×10^-2^). In addition, ETL (enzymatically treated lysate) prepared from strain N6 with an OD_600_ of 2/ml of the original culture was used. All reporter assays were performed in technical triplicates and repeated at least once independently (biological replicates) on different days. **C)** Protein quality control SDS gel showing Ni^2+^-NTA-purified recombinantly expressed heptose enzymes, GmhA, HldE (and its separate d1 and d2 domains), GmhB, all from *H. pylori* 26695a. All proteins (bands detectable at predicted masses) show a purity of >95%. HldE cloned from strain N6 was purified with similar quality features (not shown). **D)** Western immunoblot detecting purified 6xHis-HldE and separately expressed 6xHis HldE d1 and d2 domains (cloned from strain 26695a), using a custom-produced HldE antibody. The detection demonstrates that custom antibody recognizes both HldE domains and the full-length HldE protein (1:20.000, rabbit). The three main bands detected are full-length HldE (w, 52 kDa), its N-terminal domain (d1, 36 kDa), and its C-terminal domain (d2, 15 kDa) expressed separately.

**Supplemental Table S1 (Excel): Statistics (significant differences, two-way ANOVA) for comparisons of all experimental conditions performed for and depicted in Fig. 1C to 1F**.

**Supplemental Table S2 (Excel): *H. pylori* transcriptome results for all genes, analyzed for differential expression, from the transcriptomes / RNA-Seq of *H. pylori* strain Su2 (comparison between Su2 Δ*cag*PAI mutant vs Su2 wild type (wt), reference).**

