## Supplemental Figures for "*Helicobacter pylori* modulates heptose metabolite biosynthesis and heptose-dependent innate immune host cell activation by multiple mechanisms"

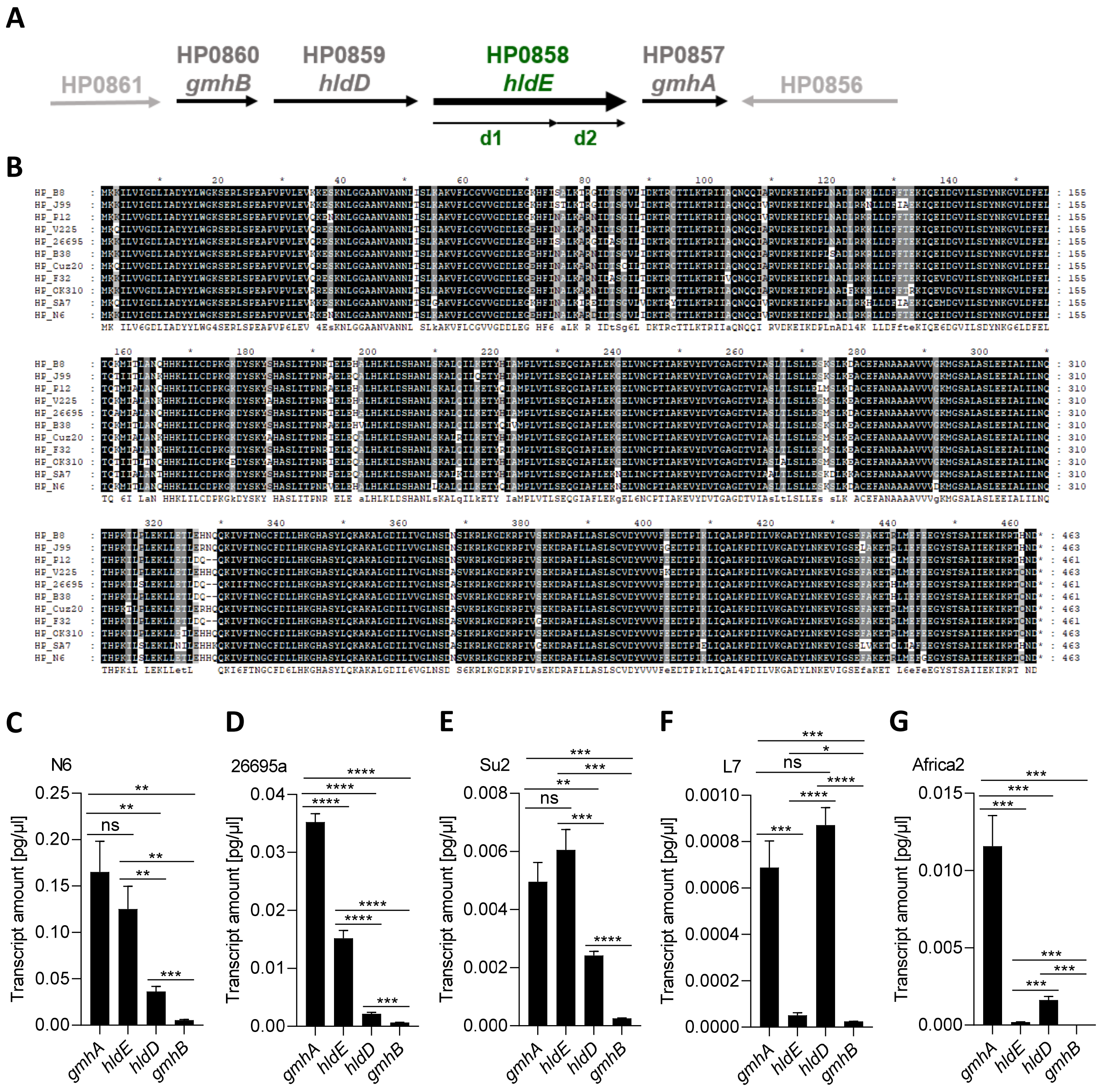


**Fig S1. Genomic arrangement, protein diversity and strain-specific regulation in the *H. pylori* heptose biosynthesis gene cluster. A**) Organisation of heptose biosynthesis operon genes HP0861 to HP0857/*gmhA* in *H. pylori* 26695a. *hldE* is highlighted in green color. Gene arrangement in the cluster is conserved in other strains isolated worldwide. **B)** variation of HP0858/HldE protein sequence in diverse *H. pylori* strains (B8, J99, V225, 26695, B38, Cuz20, F32, OK310, SA7, HPN6), selected from different geographic origins and strain populations. Note the very conspicuous hinge region in HldE between the two domains d1 and d2, characterized by gaps in some strains. **C)** to **G)** depict quantification of transcript amounts of heptose biosynthesis cluster genes *gmhA, hldE, hldD,* and *gmhB* of *H. pylori* wild type strains N6 (C), 26695a (D), Su2 (E), L7 (F) and Africa2 (G) by qPCR, performed in technical triplicates. All qPCR results, given in absolute quantities of pg/ml, were normalized to 16S rRNA transcript amounts of each sample. Pairwise significance of differences (p values) in panels C) through G) was calculated by unpaired student's *t*-test. Significance values: * $p < 0.05$; ** $p < 0.01$; *** $p < 0.001$; **** $p < 0.0001$; ns = non-significant.

**Hauke et al., Fig. S1**

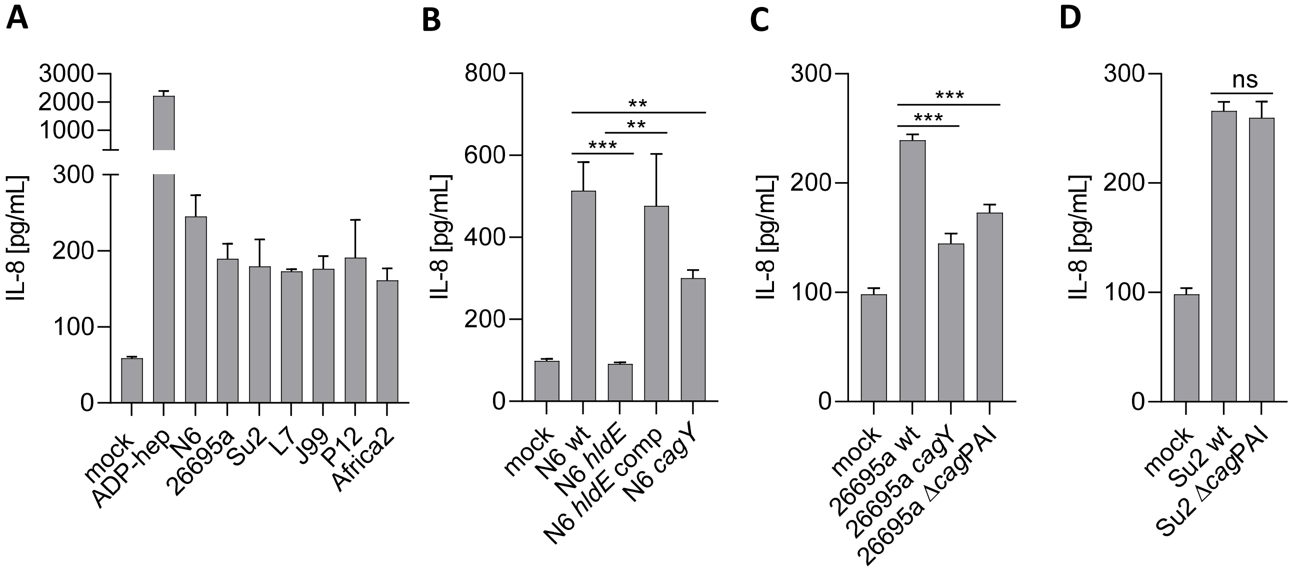


**Fig S2. Gastric epithelial cell line MKN28 response to *H. pylori* ETLs. A) to D)** Activation of MKN28 gastric epithelial cells after co-incubation with ETLs produced from various *H. pylori* wild type (wt) strains (A) and mutants (B-D) as indicated on the x-axis, for 4 h. A quantitative read-out for pro-inflammatory response was obtained by performing IL-8 ELISA. Shown are the results from technical triplicates of biological duplicates. All experiments were repeated at least once on two different days, with similar results. For B), C) and D), statistical significance was calculated for differences between wt strain and each mutant or complemented strain. Significance of differences (p values) was calculated by unpaired student's *t*-test. Significance values: ** $p < 0.01$; *** $p < 0.001$; ns = non-significant. In all cell activation experiments, co-incubation with pure ADP-heptose (2.5 µM, shown in A)) served as a reference for activation.

**Hauke et al., Fig. S2**

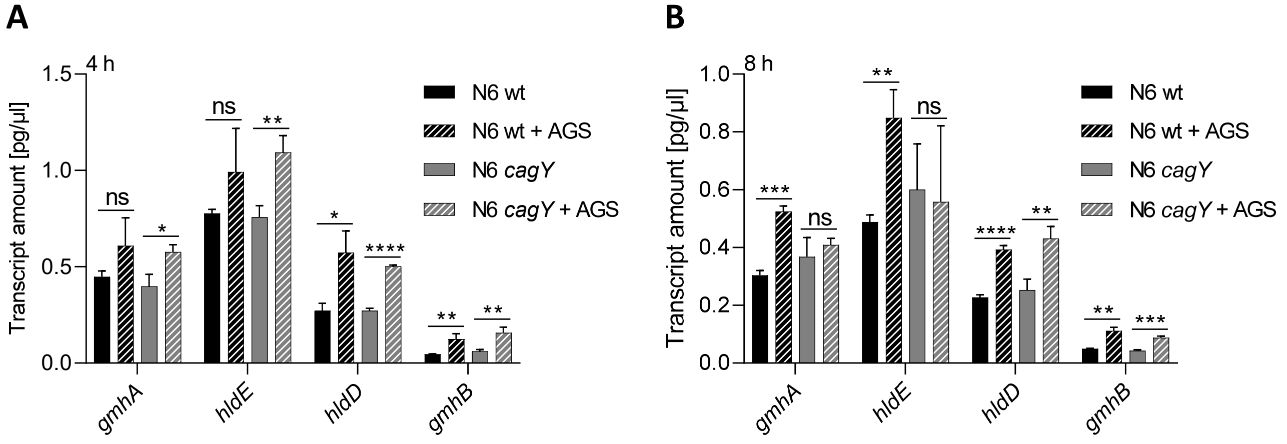


**Fig S3. Time-dependent regulation of heptose biosynthesis gene cluster in *H. pylori* N6 and its isogenic *cagY* mutant, co-incubated with gastric epithelial AGS cells. A)** and **B)** show transcript amounts (RT-qPCR) of heptose cluster genes *gmhA*, *hldE* (HP0858), *hldD*, *gmhB* (HP0860) of *H. pylori* N6 wild type and *cagY* (HP0527) mutant, both co-incubated in the presence or absence of AGS cells (MOI=50) for 4 h (A) or 8 h (B), respectively. Control bacteria were incubated in cell culture medium alone for the respective time periods. Three technical replicates are summarized in the panels. All qPCR results, shown in absolute transcript amounts of pg/µl, were normalized to 16S rRNA transcript amounts of each sample. Statistically significant differences (p) between conditions were calculated by unpaired student‘s *t*-test. Significance values: * $p < 0.05$; ** $p < 0.01$; *** $p < 0.001$; **** $p < 0.0001$; ns = non-significant.

**Hauke et al., Fig. S3**

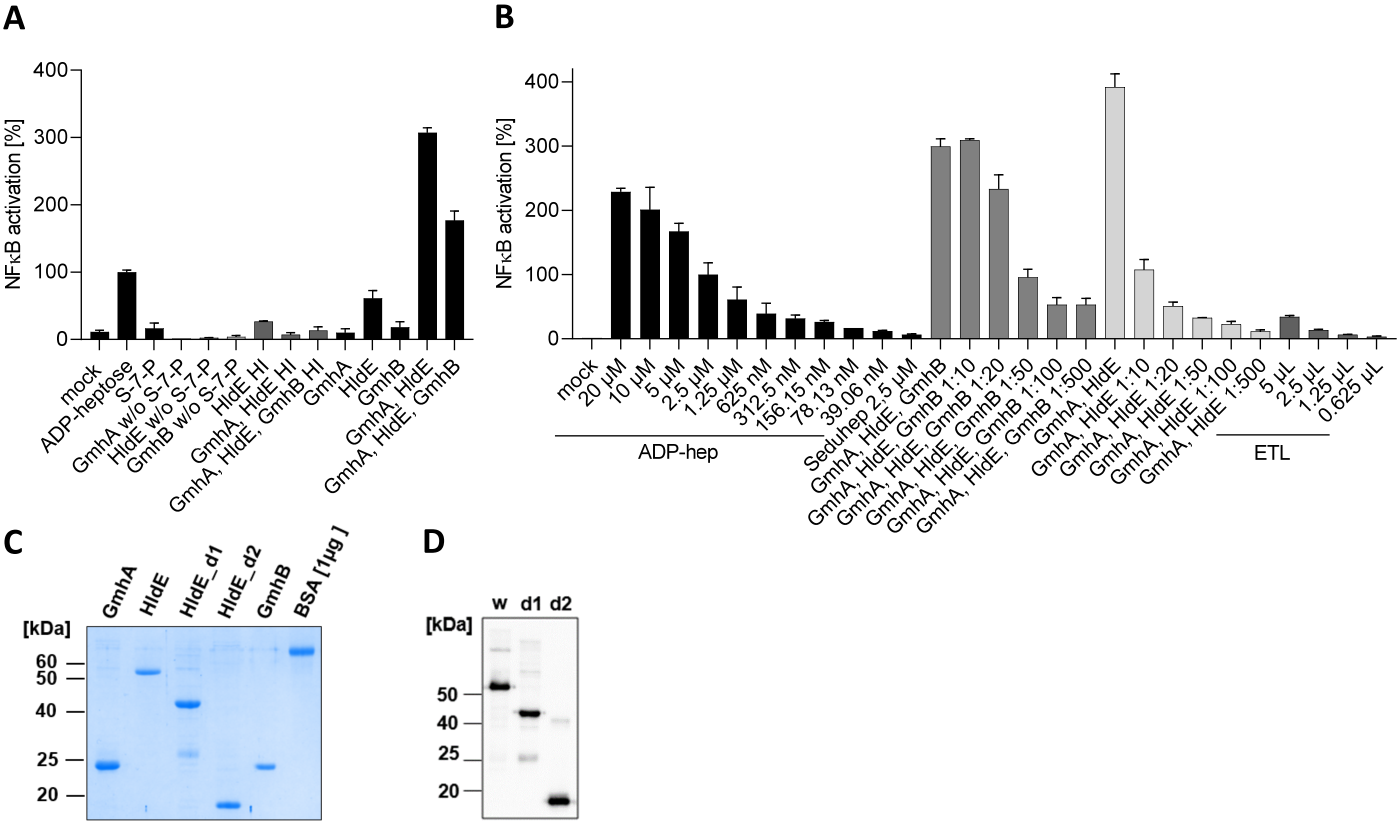


**Fig S4. Quality controls of purified recombinant *H. pylori* heptose biosynthesis enzymes and activity titrations for in vitro reconstitution of heptose biosynthesis pathway.**

**A)** Activation of NF-κB reporter cells (HEK_luc) by heptose metabolites synthesized using one-pot reactions of recombinant *H. pylori* 26695a proteins with enzymatic substrate seduheptulose 7-phosphate (S-7-P), and respective controls. Controls include active single enzymes plus substrate, or heat-inactivated (HI) enzymes plus substrate. **B)** Activation of – HEK NF-κB luciferase reporter cells by pure β-D-ADP-heptose (titration), the products of three-enzyme combination GmhA, HldE and GmhB, the reaction product of two-enzyme combination GmhA, HldE (*in vitro* reconstitution: in each case non-diluted reaction sample and the same used at different dilutions from $10^{-1}$ to $5x10^{-2}$). In addition, ETL (enzymatically treated lysate) prepared from strain N6 with an $OD_{600}$ of 2/ml of the original culture, was used. All reporter assays were performed in technical triplicates and repeated at least once independently (biological replicates) on different days. **C)** Protein quality control SDS gel showing $Ni^{2+}$-NTA-purified recombinantly expressed heptose biosynthesis enzymes, GmhA, HldE (and its separate d1 and d2 domains), GmhB, all from *H. pylori* 26695a. All proteins (bands detectable at predicted masses) show a purity of >95%. HldE cloned from strain N6 was purified with similar quality features (not shown). **D)** Western immunoblot detecting purified 6xHis-HldE and separately expressed 6xHis HldE d1 and d2 domains (cloned from strain 26695a), using a custom-produced HldE antibody. The immunoblot detection demonstrates that the custom-produced antibody (1:20.000, rabbit) recognizes the full-length HldE protein and both separate HldE domains. The three main bands detected are full-length HldE (w, 52 kDa), its N-terminal domain (d1, 36 kDa), and its C-terminal domain (d2, 15 kDa), expressed separately.
